# Particular Amino Acid on Lateral Interface Maintains FtsZ Function Under Acid Stress in *Streptococcus mutans*

**DOI:** 10.1101/2025.01.26.634952

**Authors:** Yuxing Chen, Yongliang Li, Jiahao Niu, Liuchang Yang, Yaqi Chi, Xue Cai, Fengjiao Xin, Jie Zhang, Xianyang Fang, Manas Mondal, Yiqin Gao, Xiaoyan Wang

**Author notes:** Y.C. and Y.L. contributed equally to this work. **Author Contributions:** Yuxing Chen, Yongliang Li and Xiaoyan Wang designed research; Yuxing Chen, Yongliang Li, Jiahao Niu, Yaqi Chi, Xue Cai, Jie Zhang and Manas Mondal performed the research; Yuxing Chen, Yongliang Li, Fengjiao Xin, Xianyang Fang, Manas Mondal, Yiqin Gao and Xiaoyan Wang analyzed the data; Yuxing Chen, Yongliang Li, Manas Mondal and Xiaoyan Wang wrote the paper. **Competing Interest Statement:** The authors have no competing financial interest to declare.

## Abstract

FtsZ is the core protein for cell division in bacteria that can polymerize into Z-rings and drive cytokinesis. Understanding how bacteria maintain the correct function of FtsZ under various environmental stresses is crucial for novel antibacterial drug discovery. Our previous study revealed that the FtsZ in *S. mutans* has higher self-assembly and GTPase activity under acidic stress, which may be responsible for the cariogenesis of *S. mutans*. However, the mechanism is still unknown. Here, we further reported the crystal structure of *S. mutans* FtsZ, revealing a unique lateral interface. Through protein polymerization and GTPase ability assay, we experimentally demonstrated that mutation of Arg68 on this lateral interface significantly reduced the functional activity of FtsZ in an acidic environment. The phenotype assay and rat caries model further showed that mutation of Arg68 effectively inhibited the acid resistance of *S. mutans* and the occurrence and progress of dental caries in vivo. By employing a molecular dynamics simulation analysis, we conclude that mutation of Arg68 disrupts the conformation change necessary for SmFtsZ polymerization under acidic conditions. Our study proposes a novel mechanism to maintain FtsZ function in bacteria and could be a potential target for antimicrobial drugs to inhibit the growth of *S. mutans* in an acidic environment.

**Significance Statement:** FtsZ is the core protein that drives cytokinesis. Maintaining the self-assembly and GTPase activity of FtsZ plays a critical role in cell growth under environmental stress. Currently, research on FtsZ function maintenance under stress mainly focuses on model organisms, with less on other pathogenic bacteria. This study reveals a novel structural mechanism for the acidic tolerance of FtsZ in *S. mutans* which survives in strongly acidic surroundings. Meanwhile, Arg68 on the lateral interface of SmFtsZ may be a potential therapeutic target to regulate microecology and combat *S. mutans*-associated dental caries.

## Introduction

Cell division is essential to all living organisms (1). In bacteria, numerous proteins are involved in this process, forming a complex known as the divisome (2). At its core is FtsZ, a bacterial homolog of the eukaryotic cytoskeletal protein tubulin (3–5). During cell division, FtsZ hydrolyzes GTP, polymerizes, and self-assembles into a dynamic structure known as the “Z-ring” (6, 7). This structure serves as a scaffold for the recruitment of proteins involved in peptidoglycan synthesis and chromosome segregation, ultimately facilitating cell division (2).

The self-assembly and GTPase activity of FtsZ is influenced by various environmental stresses (8, 9). Changes in ion concentrations can affect its polymerization, divalent cations enhance protofilament interactions, calcium ions promote sheet-like structures, and high magnesium concentrations (10 μM) induce the formation of elongated filament bundles (10, 11). Additionally, studies have shown that the GTPase activity of FtsZ from *Escherichia coli* (*E. coli*) and Methanococcus jannaschii significantly decreases as pH drops from neutral (7.0) to acidic (6.0), with activity becoming undetectable below pH 5.5 (8).

Several accessory proteins are recruited to the divisome, playing critical roles in cell growth under specific environmental conditions. For instance, ClpXP, a two-component protease complex, degrades polymerized FtsZ, reducing polymer abundance and altering FtsZ dynamic exchange in the Z-ring (12, 13). ClpB, a chaperone protein, prevents the accumulation of abnormal FtsZ aggregates, maintaining proper cell division under mild thermal stress. Similarly, E. coli Hsp90, a stress-responsive protein, regulates FtsZ assembly and disassembly by binding to FtsZ, either hindering its polymerization or competing with other FtsZ-interacting proteins (14). Another protein, ZapE, a member of the AAA+ ATPase family, influences the late stages of cell division. ZapE modulates FtsZ polymerization in vitro and is essential for E. coli growth under anaerobic and thermal stress conditions (15, 16).

Considering the vast differences between different bacteria, and the current results derived from model organisms, it is possible that these mechanisms are not applicable to all bacteria, especially in pathogenic species living in extreme environments. *Streptococcus mutans* (*S. mutans*), a conditional pathogen of the oral cavity, is the primary cause of dental caries (17, 18). *S. mutans* can metabolize a wide variety of carbohydrates, producing organic acids that lower the local pH and contribute to caries development. To survive this acidic environment, *S. mutans* activates an acid tolerance response, a robust mechanism that helps the bacteria adapt by buffering the cytoplasm and modifying the fatty acid composition of its membrane (19, 20). Despite these regulatory means, *S. mutans* cannot fully maintain a stable intracellular pH; its cytoplasmic pH decreases along with environmental acidification (21). However, how the function of FtsZ is regulated under the low pH condition is still unknown. Preliminary studies from our group have shown that *S. mutans* FtsZ (SmFtsZ) exhibits high acidic tolerance in vitro. It exhibits higher GTPase and polymerization activity at pH 6.0 compared to neutral pH (7.4) and retains activity even at pH 5.0 (22). In this study, we explore the mechanism behind the acidic tolerance of SmFtsZ. We report the crystal structure of SmFtsZ and highlight a unique lateral polymerization interface. We found that a specific amino acid, Arg68, located at this lateral interface, significantly affects the function of SmFtsZ under acid stress both in vitro and in vivo. Additionally, strains of *S. mutans* carrying the mutation of Arg68 (R68A) showed reduced growth under acidic conditions and cariogenesis potential. Using all-atom molecular dynamics simulations, we concluded that the R68A mutation primarily disrupts atomic interactions at the lateral interface, which are crucial for forming favorable binding interactions and maintaining the correct orientation of SmFtsZ necessary for polymer assembly under acidic conditions.

## Results

### The crystal structure of SmFtsZ exhibits a unique interface between tetramers

To investigate the mechanism underlying the acidic tolerance of SmFtsZ, we determined its crystal structure at a resolution of 3.49 Å. The crystals were assigned to the space group P4_3_2_1_2, with four monomers present in the asymmetric unit (Fig. 1*A*). Following refinement, including the addition of water molecules and metal ions, the R_work_ and R_free_ values were 25.75% and 28.74%, respectively (Table S1). The FtsZ monomer consists of two globular domains at the N- and C-terminal ends, connected by a central H7 helix (Fig. 1*B*). The N-terminal domain (residues 1–179, blue) (Fig. 1B) comprises six α-helices (H1–H6) and six β-strands (S1–S6). The C-terminal domain, shown in green, contains three α-helices (H8–H10) and four β-strands (S7– S9). Notably, the flexible T3 and T7 loops are positioned on the side and bottom surfaces of the monomer, respectively. The N-terminal domain adopts a β-α-β-α-β Rossmann fold, a characteristic feature of protein-nucleotide binding domains, which forms the GTP-binding pocket. GTP bound to SmFtsZ is shown as a rod-like structure, with Gly22, Gly23, Asn26, Ser110, Gly111, Arg144, Asn167, Leu184, and Asp188 involved in GTP binding. Structural comparisons between the crystal structure and AlphaFold3 predictions revealed a high degree of similarity in the monomeric structure, with a root-mean-square deviation (RMSD) of 0.790 Å across 285 atoms (Fig. S1*A*). The GTP-binding sites were largely conserved (Fig. S1*B*).

**Figure 1.**
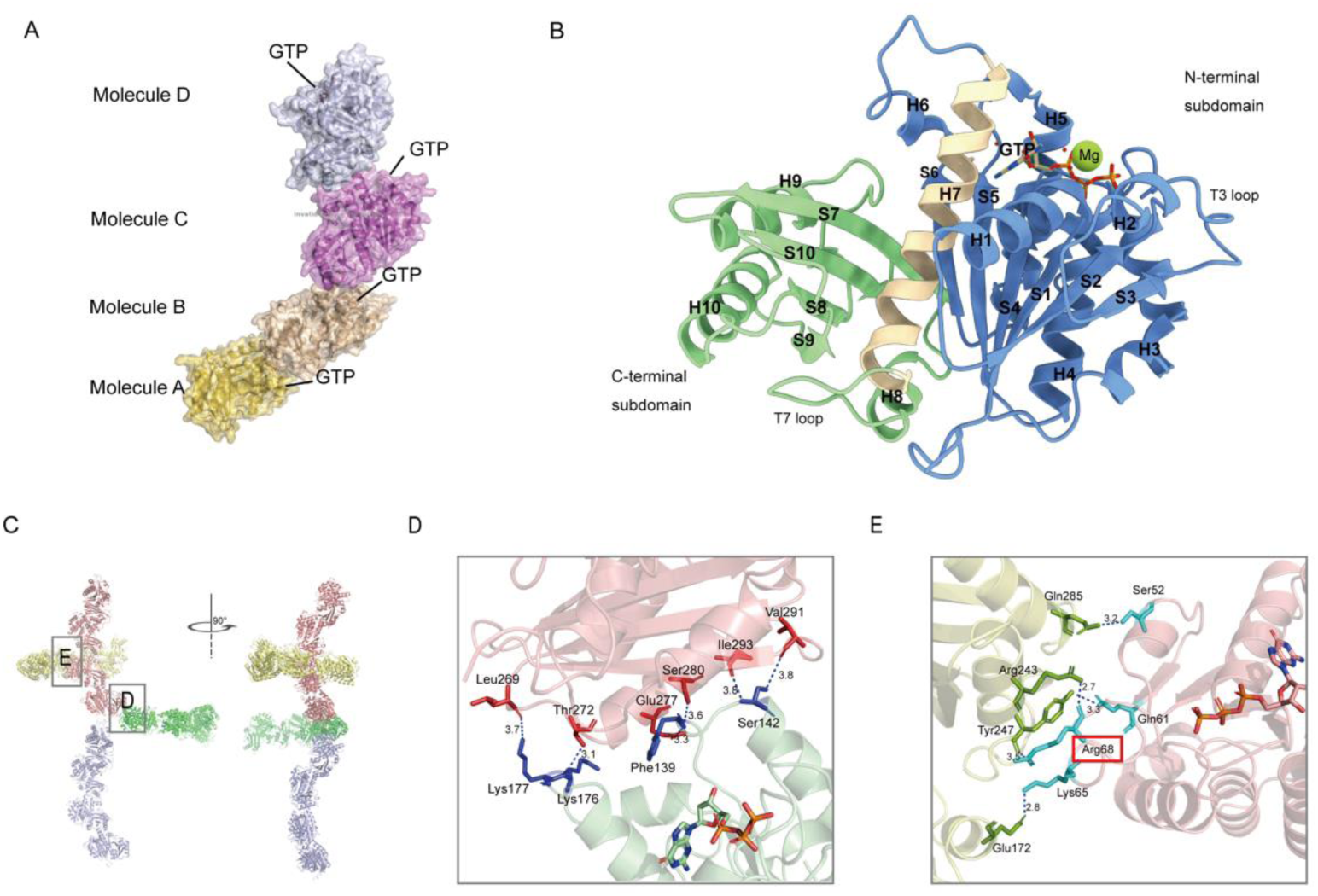
(*A*) Structure of SmFtsZ tetramers. (*B*) Structure of a single SmFtsZ monomer. (*C*) The stacking pattern of *SmFtsZ* shows a fence-like arrangement, where intertwined tetramers serve as the fundamental units. (*D, E*) Hydrogen bonds and salt bridges are formed at the lateral and longitude interfaces. The distance between two amino acids, as determined by PISA analysis, is indicated along the dotted lines.

However, the crystal structure exhibits a unique tetrameric arrangement that was not predicted by AlphaFold3 (Fig. S1*C*). The stacking pattern of SmFtsZ formed a fence-like structure, generated by the interwoven arrangement of tetramer units when the SmFtsZ tetramers are expanded (Fig. 1*C*). The longitudinal interaction interface is located between monomers and tetramers, where protein molecules connect at their head-to-tail regions. Key intermolecular interactions between amino acids at this interface include Thr272-Lys176, Glu277-Phe139, Ser280-Phe139, and others (Fig. 1*D*).

In addition, we identified a lateral interaction surface centered around the axis, characterized by five hydrogen bonds and two salt bridges. The side chains of Arg68 and Lys65 in the T3 loop form hydrogen bonds with Arg243/Tyr247 and Glu172, respectively. Further hydrogen bonds were observed between Gln61 and Tyr247 and Ser52 and Gln285 (Fig. 1*E*).

### R68 is essential for FtsZ polymerization in acidic environments in vitro

To determine whether intermolecular forces at the lateral interface influence FtsZ polymerization, we introduced mutations at five side-chain amino acid positions selected to minimize disruption to overall protein folding. The filament structures of the resulting variants were examined under varying pH conditions using transmission electron microscopy (TEM). Furthermore, side-chain lengths were varied at specific amino acid positions to assess their effects on polymerization.

Among the five amino acid residues tested, E277 mutations had no significant impact on SmFtsZ polymerization. Variants SmFtsZ-E277T, SmFtsZ-E277L, and SmFtsZ-E277D all formed longer protofilaments at pH 7.4 and assembled into multi-filament bundles at pH 6.0 and 5.0, similar to the wild-type SmFtsZ (SmFtsZ-WT) (Fig. 2*A* and Fig. S2). By contrast, mutations at T272 and S280 severely disrupted polymerization. SmFtsZ-T272V, SmFtsZ-T272C, SmFtsZ-S280A, and SmFtsZ-S280R all failed to assemble into filaments even under pH 7.4 condition (Fig. 2*B* and Fig. S2).

**Figure 2.**
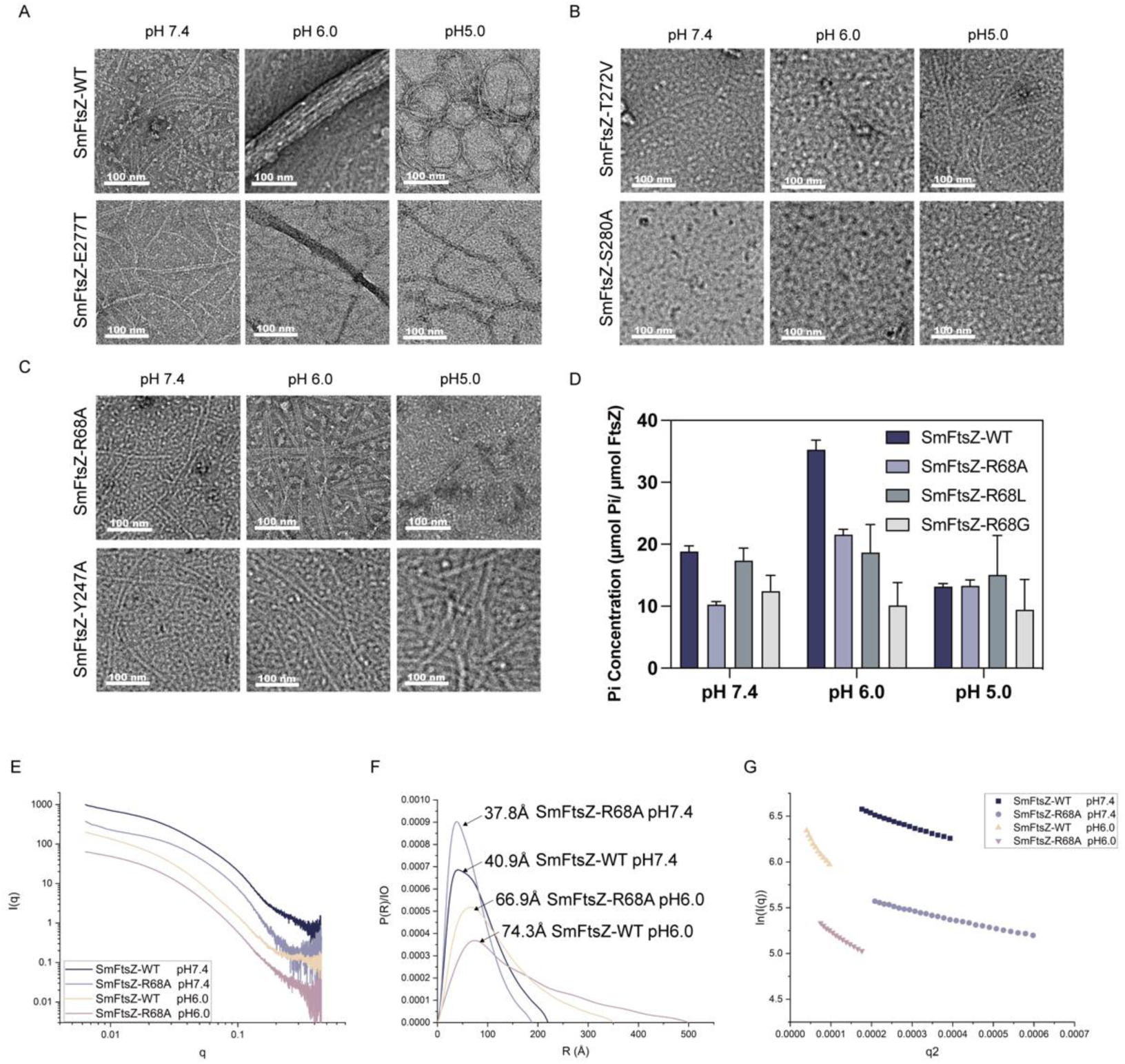
Effect of pH on the GTPase activity and polymerization of SmFtsZ-WT and its variants. (*A–C*) The effect of pH on the polymerization of SmFtsZ-WT and its mutations was examined. SmFtsZ (10 µM) was added to polymerization buffer at pH 7.4, 6.0, and 5.0 and incubated for 15 min with 2 mM GTP. For negative staining, samples were exposed to a 1% solution of uranyl acetate. Scale bar: 100 nm. (*D*) GTPase activity of *SmFtsZ-WT* and the mutants *SmFtsZ-R68A*, *SmFtsZ-R68L*, and *SmFtsZ-R68G* at different pH values. (*E–G*) SAXS analysis of SmFtsZ-WT and SmFtsZ-R68A under varying pH conditions. The scattering profiles (*E*), pair distance distribution functions (PDDFs) (*F*), and Guinier fitting plots (*G*) of SmFtsZ-WT and SmFtsZ-R68A at pH 7.4 and pH 6.0, respectively.

Interestingly, we found that SmFtsZ-R68A and SmFtsZ-Y247A both formed filament structures similar to those of SmFtsZ-WT at pH 7.4. However, unlike the SmFtsZ-WT, these mutations were unable to form multi-filament bundles or cyclic polymers at pH 6.0 and 5.0, indicating that R68 and Y247 significantly influence polymerization under acidic conditions (Fig. 2*C*). But SmFtsZ-Y247F exhibited filament structures comparable to those of the SmFtsZ-WT in acid environments, suggesting that the effect of Y247 on polymerization may depend on the side-chain length. Notably, consistent with SmFtsZ-R68A, SmFtsZ-R68G and SmFtsZ-R68L also impaired SmFtsZ polymerization under acidic conditions, suggesting that R68 is a critical site influencing the polymerization of SmFtsZ under acidic conditions.

Additionally, we also observed that the GTPase activity of SmFtsZ-R68 mutants was reduced to varying degrees compared to SmFtsZ-WT, with the most pronounced decrease at pH 6.0 (Fig. 2*D*). This reduction in enzymatic activity correlates with impaired filament assembly and lateral bundling, highlighting the importance of R68 in maintaining the stability of FtsZ polymerization in acidic environments.

Next, we employed small-angle X-ray scattering (SAXS) to further characterize the polymerization behavior of SmFtsZ-WT and SmFtsZ-R68A in solution at pH 7.4 and 6.0. The scattering patterns, represented as scattering intensity I(*q*) versus momentum transfer q, along with the pair-distance distribution functions (PDDFs) and dimensionless Kratky plots derived from these profiles for SmFtsZ-WT and SmFtsZ-R68, are shown in Fig. 2E–G. Key structural metrics, such as the radius of gyration (R_g_) and the maximum end-to-end distance (D_max_), were higher for SmFtsZ at pH 6.0 compared to pH 7.4 (Table S2). This suggests increased polymerization under acidic conditions.

The PDDFs (Fig. 2*F*) confirmed that all polymerized SmFtsZ elongated in solution. For SmFtsZ-WT, the primary distance distribution values at pH 7.4 and pH 6.0 were approximately 40.9 Å and 74.3 Å, respectively. In comparison, for SmFtsZ-R68A, the primary distance distribution values were 37.8 Å and 66.9 Å, reflecting significant differences in the cross-sectional radius. Guinier analysis of the scattering data further corroborated these findings (Fig. 2*G*). SmFtsZ-WT formed longer, thicker filament bundles under acidic conditions, consistent with increased polymerization. By contrast, SmFtsZ-R68A formed shorter, thinner filaments at pH 6.0, indicating the essential role of R68 residue in promoting lateral filament interactions and stabilizing the polymer structure under acidic stress.

### R68 is essential for FtsZ function in vivo

The treadmilling activity of FtsZ and its correct localization at the cell center is critical for its function. We assessed the treadmilling behavior of FtsZ filaments using total internal reflection fluorescence (TIRF) imaging (Fig. 3*A*). At pH 5.0, SmFtsZ-WT displayed treadmilling activity with an average speed of 36.95 ±10.57 nm/s, which was similar to that observed at pH 7.4. However, the introduction of the R68A mutation significantly affected treadmilling behavior. While treadmilling was observed at pH 7.4 with an average speed of 32.98 ± 8.964 nm/s, the speed was markedly slower at pH 5.0 (7.061 ±3.002 nm/s) (Fig. 3*B*).

**Figure 3.**
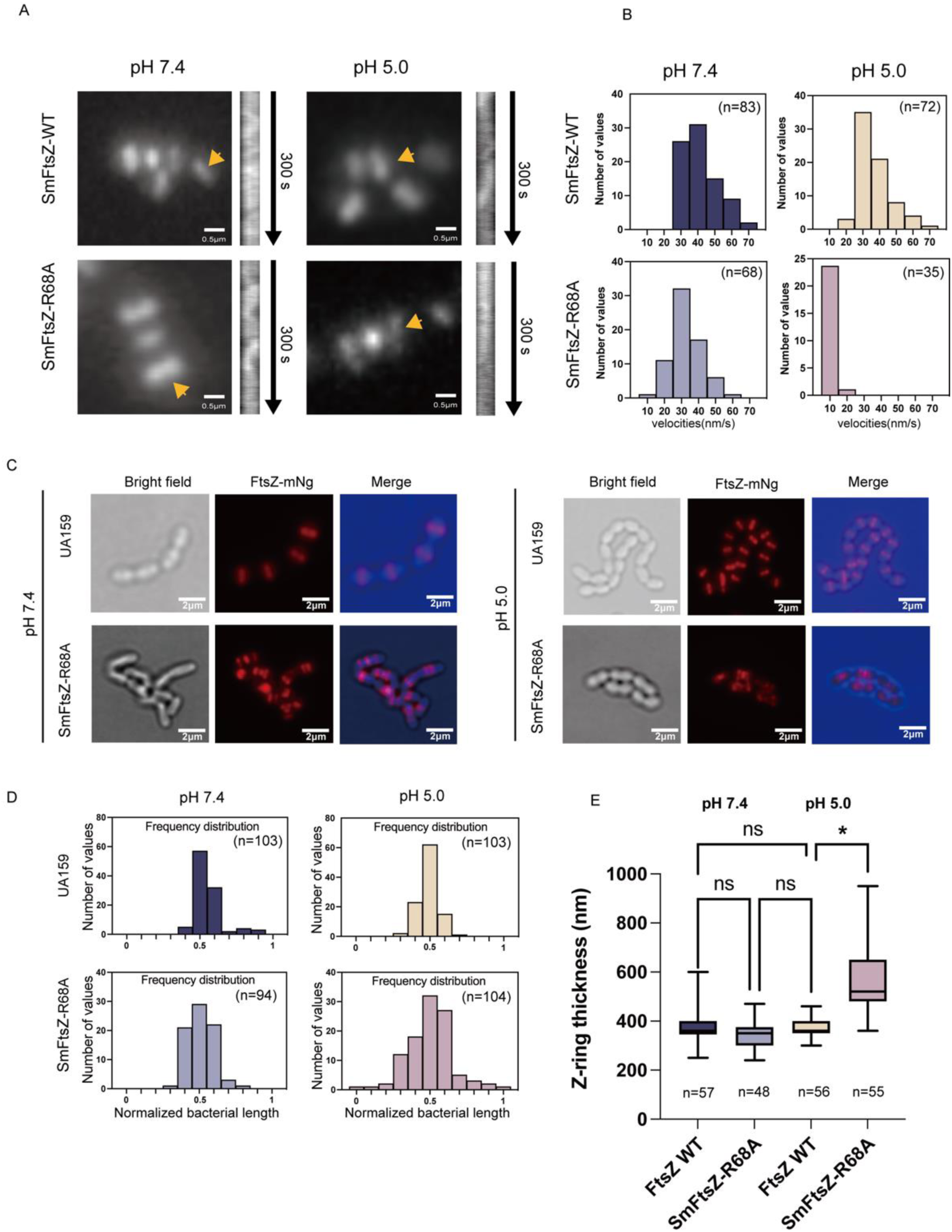
Characterization of SmFtsZ mutations in vivo. (*A*) Visualization of FtsZ ring dynamics using *FtsZ_mNeongreen*, showing the directional movement of SmFtsZ-WT and SmFtsZ-R68A rings in cells. Kymographs illustrate the dynamics of FtsZ filaments within the Z-rings (yellow arrows). (*B*) Velocity distribution of FtsZ filaments within the Z-rings of cells (n = the number of events analyzed). (*C*) Representative images of FtsZ rings in cells at pH 7.4 and 5.0 are shown. Brightfield and fluorescence microscopy were used to analyze *S. mutans* cells expressing *FtsZ_mNeongreen* (red). Scale bar: 2 μm. (*D*). Cellular distribution of FtsZ. Bacterial length is normalized to 1, with 0 and 1 on the X-axis representing the two poles of the cell, and 0.5 indicating the midpoint of the cell (n = the number of cells analyzed). (*E*) Z-ring thickness for SmFtsZ-WT and SmFtsZ-R68A under different pH conditions. Statistical assessments were conducted using one-way ANOVA with GraphPad Prism 9 (GraphPad Software, La Jolla, CA, USA). ‘*’ indicates *p* < 0.05. ‘ns’ indicates no significant difference, and n represents the number of cells analyzed.

Next, we investigated the spatial localization of SmFtsZ-R68A. SmFtsZ-WT localized correctly to the center of the bacterial cell, regardless of whether cells were grown in the medium at pH 7.4 or pH 5.0 (Fig. 3*C*). By contrast, SmFtsZ-R68A predominantly localized to the cell center or future division site in pH 7.4 medium. However, under acidic conditions (pH 5.0), localization was disrupted, with some failing to localize to the division site properly (Fig. 3*D*). Furthermore, we assessed Z-ring thickness and found that SmFtsZ-R68A and SmFtsZ-WT showed similar Z-ring thickness at pH 7.4. However, SmFtsZ-R68A exhibited a thicker ring structure at pH 5.0 than that of SmFtsZ-WT, indicating that SmFtsZ-R68A formed a more loosely packed Z-ring at pH 5.0 (Fig. 3*E*).

### R68 mutation impaired the growth of *S. mutans* in acidic environments

To determine whether the R68A-mediated dysfunction of FtsZ affects the pathogenicity of *S. mutans*, we examined the growth of the SmFtsZ-R68A strain under various pH conditions. At pH 7.4 and 6.0, the R68A mutation produced no significant impact on the growth of *S. mutans* (Fig. 4*A*). However, at pH 5.0, the R68A strain exhibited a substantial reduction in growth rate and overall biomass. Furthermore, more dead cells were detected in the SmFtsZ-R68A strain compared to the SmFtsZ-WT strain in 96 hours biofilms (Fig. 4*B-C*), indicating a marked reduction in acid tolerance for the SmFtsZ-R68A strain.

**Figure 4.**
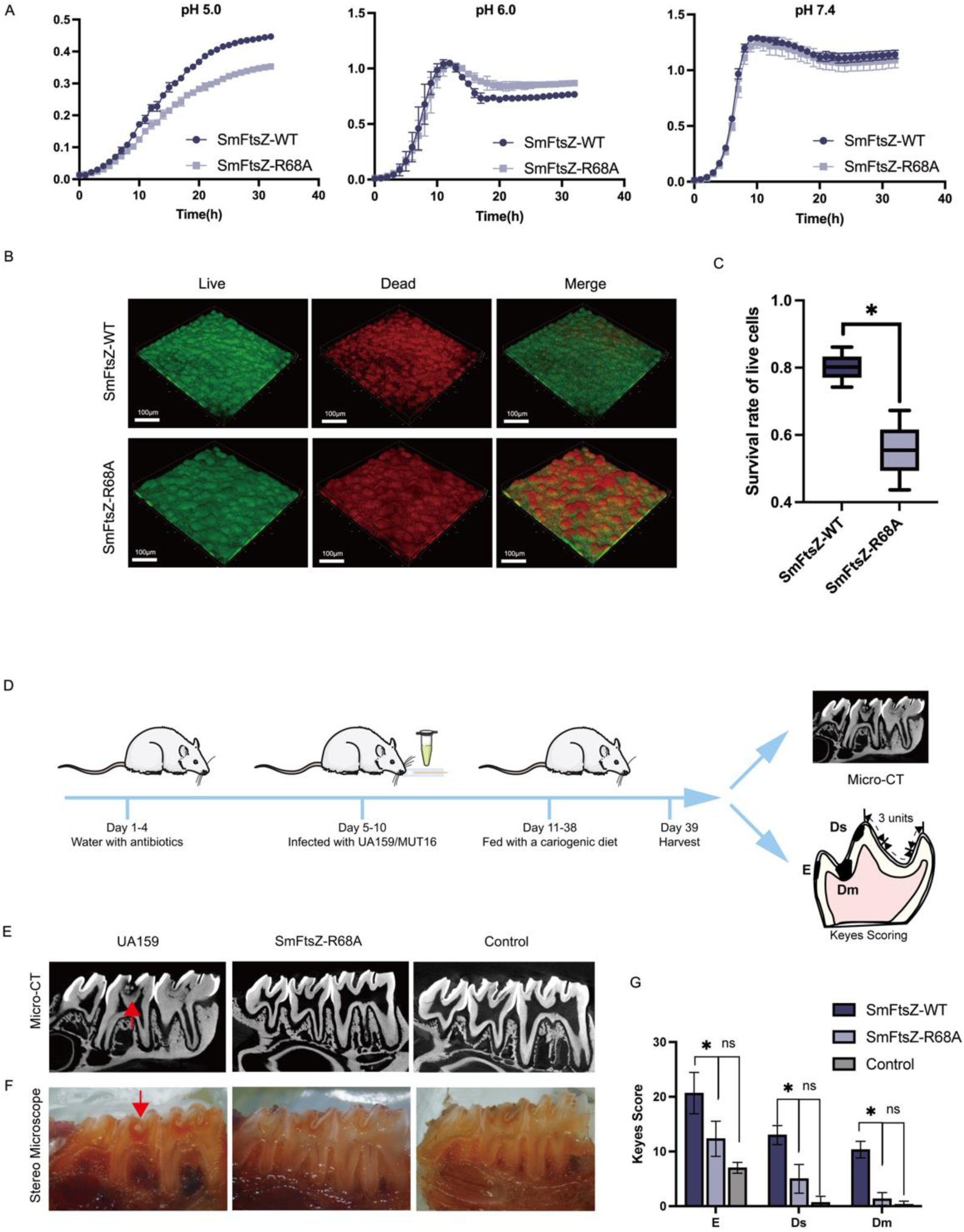
Effects of acidic conditions on *S. mutans* UA159 and SmFtsZ-R68A strains. (*A*) Growth curves of *S. mutans* UA159 strain and SmFtsZ-R68A strains at different pH levels. Data were collected from three separate and independent experiments. (*B*) Representative CLSM images of biofilms of *S. mutans* UA159 and SmFtsZ-R68A strains. Live cells (green) were stained with SYTO 9, while dead cells (red) were stained with propidium iodide. Images were captured using a 25× objective lens. Scale bar: 100 µm. (*C*) Survival rate of live cells of biofilm analysis using Leica imaging software. Data represent the means of three independent experiments, analyzed statistically with the Mann−Whitney U test. ‘*’ indicates *p* < 0.05. (*D*) Overview of the surgical procedure for establishing a dental caries model. (*E–F*) Representative micro-CT and stereomicroscope images of molar regions in the dental caries model are shown in sagittal view. (*G*) Keyes’ scores for the three groups representing levels of dental cavities: Dm refers to moderate dentinal, Ds indicates slight dentinal, and E represents enamel. Data were analyzed using one-way ANOVA in GraphPad Prism 9 (Statistical analysis was performed using one-way ANOVA with GraphPad Prism 9 (GraphPad Software, La Jolla, CA, USA). ‘*’ denotes *p* < 0.05, while ‘ns’ represents no significant difference.

Next, we established an in vivo rat dental caries model. The surgical procedures are outlined in Fig. 4*D*. No bacterial colonies were detected in the rats after streptomycin treatment. However, colonies of *S. mutans* and the SmFtsZ-R68A strain were successfully cultured following oral inoculation (Fig. S3). Throughout the study, rat body weight remained stable across all experimental groups. Representative micro-CT and stereomicroscope images of the molar regions in sagittal view are shown in Fig. 4 *E-F*. While varying levels of carious lesions were observed in the UA159 group (SmFtsZ-WT), dentin lesions were rarely detected in the SmFtsZ-R68A and control groups. The Keyes’ scoring system, a widely used method for evaluating caries, categorizes four types of pit and fissure lesions based on depth: enamel only (E), slight dentinal (Ds), and moderate dentinal (Dm). The total Keyes’ scores for all lesion types were markedly reduced in the SmFtsZ-R68A strain compared to the SmFtsZ-WT (E: *p*<0.05, Ds: *p*<0.05, Dm: *p*<0.05) (Fig. 4*G*). These findings show that the SmFtsZ-R68A mutation effectively reduces the onset and progression of dentin caries in the rat model.

To assess whether the effects of SmFtsZ-R68A on acidic resistance were independent of known regulatory systems, we investigated changes in RNA translation levels in the SmFtsZ-R68A and UA159 strains. Differential expression analysis revealed 268 upregulated and 318 downregulated genes between the two strains. However, genes involved in key acid tolerance mechanisms, including F1F0-ATPase, the agmatine deiminase system (AgDS), and membrane fatty acid composition, showed no significant changes (Table. S3).

### Insights into intermolecular interactions at the binding interface of SmFtsZ in acidic pH environments from MDS

Biophysical studies have characterized how acidic pH and mutations influence the GTPase activity and polymerization of SmFtsZ. To further understand the atomistic interactions at the interface of SmFtsZ assembly in a pH-dependent manner, we performed all-atom constant pH molecular dynamics (CpHMD) simulations of the SmFtsZ dimer in explicit solvent at pH 5.0, 6.0, and 7.0 (23). These simulations were conducted with longitudinal SmFtsZ-dimer_long_ and lateral SmFtsZ-dimer_late_ interaction interfaces (Fig. S5). Details of the simulation methodology are described in the Supporting Information (SI). The simulations revealed that solvent pH affects the conformation of SmFtsZ-dimer_long_ and SmFtsZ-dimer_late_ and induces residual fluctuations of the residues in the SmFtsZ chains (Fig. S6).

The average superposed structures of SmFtsZ-dimer_long_ and SmFtsZ-dimer_late_ at different pH values (Fig. S7) indicate that pH primarily alters the mutual orientation of SmFtsZ within the dimeric complex, thereby affecting interactions at the binding interface. A major change in mutual orientation was observed in SmFtsZ-dimer_late_ at pH 5.0. Residue pair distance distributions in the protein chains at different pH values (Fig. S8), which primarily form contacts in the dimeric interface as seen in the initial crystal structure, showed that pH variations alter key protein contacts and interactions within the binding interface for SmFtsZ-dimer_long_ and SmFtsZ-dimer_late_ complexes. In the longitudinal dimeric interaction interface, the residue pairs Thr272-Lys176, Glu277-Phe139, Ser280-Phe139, and Ile293-Ser124 maintained favorable contacts at pH 6.0 (Fig. S8*A–D*). In the lateral dimeric interface, the residue pairs Lys65-Glu172, Arg68-Tyr247, and Gln61-Tyr247 maintained favorable contacts at pH 6 and 7 (Fig. S8*E-H*). An acidic pH environment alters the interface area and binding interactions within the dimeric complex (Fig. S9, see the SI for details).

We further analyzed the protonation states of the acidic residues and potential hydrogen bonding interactions within the dimeric interface at different pH values. The side chains of most of the Asp and Glu residues near the dimeric interface exhibited higher protonation probabilities at pH 5.0, with a subset also protonated at pH 6.0 (Fig. S10). Thus, environmental pH affects the protonation states of acidic residues at the binding interface, which modulates the binding interactions between SmFtsZ monomers at the longitudinal and lateral interaction interfaces. The interactions among the key interface residues in SmFtsZ-dimer_long_ (Fig. S11) are described in detail in the SI.

Hydrogen-bonded interactions and the orientation of key interface residues in the lateral interaction interface SmFtsZ-dimer_late_ were examined at different pH values (Fig. 5*A–C*). At pH 7.0, the side chain of Arg243 in chain B and Arg68 in chain A form hydrogen-bonded contacts with the backbone oxygen atom of Arg68 in chain A and Tyr247 in chain B, respectively. The side chain of Arg68 and Lys65 in the T3 loop of chain A also form hydrogen-bonded contacts with Glu252 in chain B at the interface at pH 7.0 (Fig. 5*A*). At pH6.0, Arg68 in the T3 loop of chain A forms hydrogen-bonded contacts with Tyr247 and Leu251 in chain B, while Gln49 in chain A forms a hydrogen bond with the side chain of Arg-243 in chain B (Fig. 5*B*).

**Figure 5.**
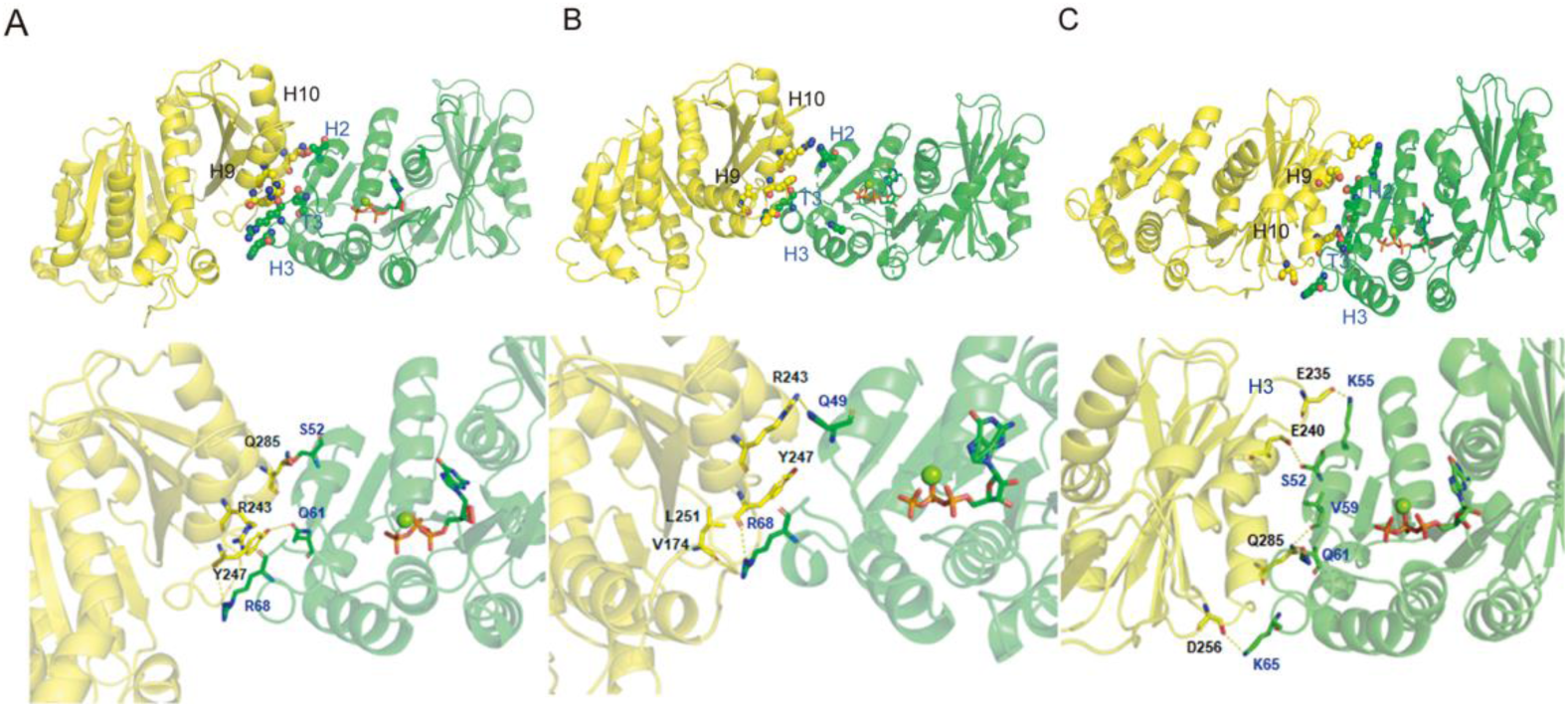
Dimeric binding interface and interface residues forming potential hydrogen bonds between the protein chains in SmFtsZ-dimerlate complexes at (*A*) pH 7.0, (*B*) pH 6.0, and (*C*) pH 5.0.

At pH 5.0, significant changes occur in the mutual orientation of protein chains at the lateral interaction interface. These changes include shifts in the orientation of the H9 helix in chain B, which contains Tyr247, resulting in the formation of new residue contact pairs within the interface because of proximity with the H2 helix of chain A (Fig. 5*C*). At this pH, the T3 loop of chain A establishes different contacts with chain B at the interface. Specifically, Ser52, Val59, and Gln61 in chain A form favorable hydrogen-bonded interactions with Glu240 and Gln285 in chain B, respectively. Furthermore, the formation of salt bridges between Lys55 and Lys65 in chain A and Glu235 and Asp256 in chain B, respectively, increased the interaction interface and stabilized the lateral interaction at pH 5.0 (Fig. 5*C*).

We calculated the free energy of binding (ΔG) between the SmFtsZ monomers in the SmFtsZ-dimer_late_ complex using the molecular mechanics generalized born surface area (MMGBSA) approach (24). The total ΔG at pH 6.0 was significantly more negative (−20.24 ±7.9 kcal/mol) than the values at pH 5.0 (−12.55 ±12.1 kcal/mol) and pH 7.0 (−8.75 ±11.6 kcal/mol), indicating a higher binding affinity between the protein monomers at pH 6.0.

Alanine scanning mutagenesis of Arg68 at the lateral dimeric interaction interface revealed that the ΔΔG value, which measures the relative effect of the R68A mutation on the ΔG of binding, becomes positive with ΔΔG ∼ 3.83 (±2.2), 9.24 (±3.9) and 3.63 (±2.5) kcal/mol at pH 7.0, 6.0 and 5.0, respectively. Thus, the R68A mutation can reduce the binding affinity between SmFtsZ monomers and significantly affect the binding interaction at pH 6.0. MDS further revealed that the R68A mutation in the SmFtsZ-dimer_late_ complex did not significantly affect the conformation of individual protein chains (Fig. S12 *A–B*). However, it significantly altered the dimeric conformation by changing the mutual orientation of the protein chains and interface contacts under acidic conditions (Fig. S12 *C–E*), compared to the native structure. The average number of potential hydrogen bonds between the protein chains decreased with the R68A mutation (Fig. S12*F*), and a significant reduction in the interaction energy was observed at pH 6 because of the mutation (Fig. S12*G–H*). The simulated average structure of the mutated dimeric complex (Fig. 6) also showed that the mutation had a differential impact on dimeric conformation at pH 6.0 and 5.0, resulting from changes in interface contacts. Interface contacts involving the H2 helix and T3 loop of chain A with chain B were largely shifted. Specific residue pairs of Arg243-Arg68Ala, Tyr247-Arg68Ala, and Tyr247-Gln61 were absent at the dimeric interface at pH 6.0, because of the large-scale movement of the H9 and H10 helices of chain B relative to the H2 helix of chain A in the mutated complex (Fig. 6). Conversely, at pH5, the H9 and H10 helices of chain B are present at the dimeric interface site in the mutated dimeric complex but adopt a different orientation, forming fewer contacts with the chain A (Fig. 6) compared to the native structure. Furthermore, because of the R68A mutation in the T3 loop, the H2 helix of chain A lost interface contacts with the H9 helix of chain B. At pH 5.0, Tyr247 and Arg135 in chain B form hydrogen bonds with Gln61 and Glu92 in chain A, respectively. These findings suggest that the R68A mutation primarily affects atomic interactions among specific residues at the lateral interface, disrupting favorable binding interactions and altering the mutual orientation of SmFtsZ monomers necessary for polymeric assembly under acidic conditions.

**Figure 6.**
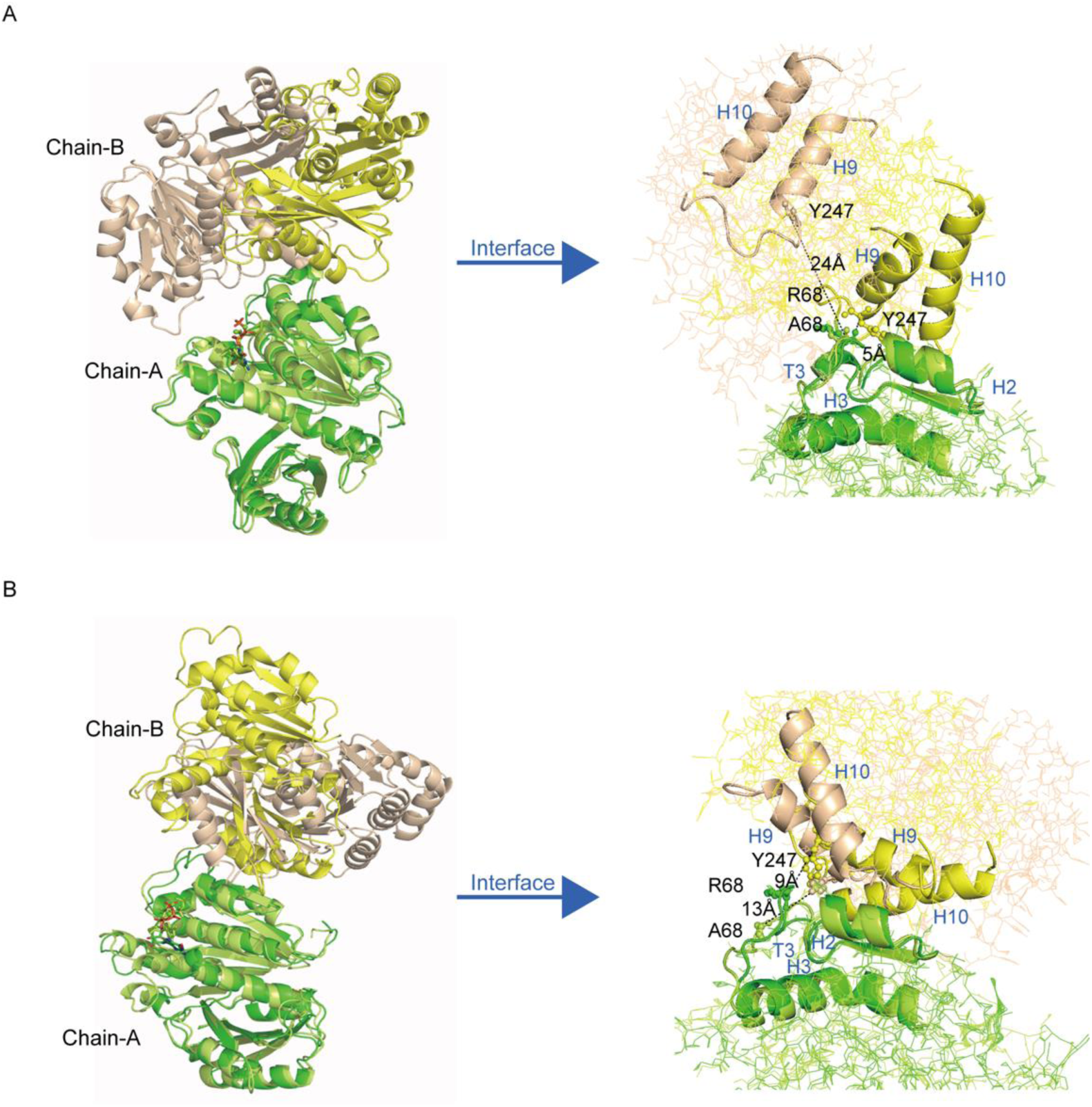
Superposed simulated average structure and interface helices of the native and R68A-mutated lateral dimeric conformation of SmFtsZ at (*A*) pH 6.0 and (*B*) pH 5.0

## Discussion

To shed light on the mechanism of acid tolerance of SmFtsZ. This study presents the first crystal structure of FtsZ from Streptococcus mutans at a resolution of 3.5 Å. The overall monomer conformation aligns with previously reported FtsZ structures and predictions from Alphafold3. Notably, SmFtsZ assembles head-to-tail into a unique tetramer structure and cross stacks into a “fence-like” pattern, a structural arrangement not previously observed in other species. This distinctive arrangement highlights novel lateral interaction interfaces involving the T3 loop in its polymerized form. Previous studies have reported the lateral interaction of FtsZ in other species, such as *Klebsiella pneumoniae* or *E. coli*. However, these studies used monobody or amino acid mutation to stabilize the filaments, which might disrupt FtsZ function in vivo, suggesting that the lateral interaction they reported is not physiological (25, 26). In contrast, the lateral interaction of SmFtsZ we observed appears closer to the physiological state.

Through PISA (http://www.ebi.ac.uk/pdbe/pisa) (27) analysis, we identified amino acid pairings that form hydrogen bonds and salt bridges at the interface, generating robust intermolecular forces. Mutations such as T272V and S280A led to a complete loss of polymerization, further validating the importance of this lateral binding interface to SmFtsZ function. Furthermore, the R68 mutation specifically disrupted FtsZ polymerization under acidic conditions but not in neutral environments, suggesting a pH-dependent role for R68 in maintaining FtsZ polymerization. We also observed decreased GTPase activity in the R68A mutant. As R68 is not involved in GTP binding sites and FtsZ polymerization is a prerequisite for GTPase activity, we infer that while R68A does not affect GTP binding, weakened FtsZ polymerization resulted in the release of fewer phosphorus ions (Fig. 2*C–D*). SAXS analysis confirmed that the SmFtsZ-R68A mutation decreased polymerization size at pH 6.0. Considering that Arg carries a positive charge, it likely forms electrostatic forces on the lateral interaction interface. This is consistent with Beuria’s observation that ionic strength affects FtsZ protofilament bundling (28).

The dynamic treadmilling of the Z-ring, vital for FtsZ function (29), has been well-documented in Escherichia coli and Bacillus subtilis (30, 31). In this study, we found that the treadmilling speed of SmFtsZ-R68A was only slightly affected at pH 7.4, but it was significantly increased at pH 5.0 (Fig. 3*B*). The treadmilling movement of FtsZ is tightly dependent on FtsZ’s GTPase activity (32). According to the “cytomotive switch” model, polymerization- and depolymerization-driven conformational changes of FtsZ are required for treadmilling of the protofilaments (33). The present study proved that mutations outside the GTP binding site could disrupt FtsZ treadmilling, indicating that impaired polymerization directly affects the motility function of FtsZ within the cell, supporting the cytomotive switch model.

Moreover, we observed that the SmFtsZ-R68A caused noticeable dispersion of FtsZ localization (Fig. 3*C*) and thicker Z-rings (Fig. 3*E*) under acidic conditions, indicative of a loose Z-ring. Two plausible explanations for this phenotype are: (a) the SmFtsZ-R68A mutation slows FtsZ treadmilling speed, which may be critical for Z-ring condensation, and (b) the R68A mutation alters lateral protofilament interaction in the Z-ring, affecting Z-ring condensation and localization.

To elucidate the mechanism by which protein conformational changes participate in protein acid resistance, we used CpHMD to investigate how pH changes alter the conformational changes in the FtsZ lateral interaction interface. These revealed distinct dimer conformations of SmFtsZ at pH 7.0, 6.0, and 5.0, driven by changes in lateral interfacial interactions (Fig. 5). We further identified the specific interactions within the two different binding interfaces that stabilize the dimeric assembly, with variations in binding interactions across different pH conditions (Fig. S6). These interactions likely contribute to the stability and functionality of FtsZ polymerization in acidic environments. The identification of favored intermolecular interactions between FtsZ, mediated by specific hydrogen bonding and salt bridge interactions at pH 6.0, aligns with our experimental observations of increased FtsZ polymerization at this pH and provides atomistic insights for this process. At pH 5.0, significant conformational changes in FtsZ dimers were observed with lateral interaction interfaces caused by altered residue interactions at the lateral interface. These changes may protect FtsZ function under acidic conditions.

We individually mutated the amino acids at the lateral interaction interfaces and tested the function of the mutant proteins in vitro. Our results showed that the Arg68 mutation significantly reduced the lateral polymerization at pH 6.0; GTPase activity was lower than that of the SmFtsZ-WT. These experimental results were consistent with structural and simulation data, confirming that Arg68 is critical for forming hydrogen bonds with Tyr247 and Arg243 at the lateral interface. The R68A mutation reduced binding interactions and disrupted the particular dimer conformation at pH 6.0 by altering interactions among key residues in the lateral dimeric interface (Fig. 6). We conclude that the polymerization morphology influences the acid resistance properties of the protein.

Although multiple bacteria contribute to dental caries, targeting *S. mutans* in dental biofilm is a practical approach to prevent caries (34). This is mainly because *S. mutans* shows an outstanding ability of biofilm formation and cariogenic (35, 36). Interestingly, our study proved the R68 mutation not only reduces the growth ability of *S. mutans* under acidic conditions and the proportion of viable bacteria in matured biofilm (Fig. 4 and Fig. S3) but also weakens *S. mutans* cariogenic ability in a rat dental caries model. Traditional anti-caries prevention strategies focus on eradicating oral bacteria and eliminating biofilms, targeting both cariogenic and commensal bacteria with broad-spectrum antibacterial drugs. However, the obvious shortcomings are that these drugs may increase bacterial antibiotic resistance and even affect the balance of oral microecology. Novel ecological antimicrobial approaches to dental caries focus on inhibiting *S. mutans* without affecting the diversity of oral microbiota, which emphasizes the crucial role of establishing a healthy microbiome in caries prevention. The findings of our study show Arg68 is identified as a potential target for inhibiting *S. mutans* growth in a low pH and cariogenic environment, instead of affecting the resistance of *S. mutans* in the natural oral microecology.

## Materials and Methods

### Bacterial Strains, Plasmids, and Media

The plasmids, amplicons, and bacterial strains used are listed in Supplementary Table S4. The *S. mutans* UA159 WT strain and its mutants were maintained on BHI media (BBL Becton Dickinson). By contrast, *Escherichia coli* Top10 strains were grown on LB media. Both media were supplemented with antibiotics (Table S4). All *S. mutans* cultures were maintained without agitation at 37 °C in an atmosphere containing 5% CO₂ and 95% air.

### Protein expression and purification

Full-length SmFtsZ and its amino acid mutations were used. The FtsZ derivatives were prepared as C-terminal His6-tagged fusion proteins using the pET28a vector. The full-length SmFtsZ expression vectors were developed in an earlier study conducted by our group (22). Site-directed SmFtsZ mutations were introduced using a quick-change replacement strategy (see Table S5 for the primers used). The mutations were confirmed via sequencing. For protein expression, plasmids were introduced into *E. coli* BL21 (DE3). The procedure for expressing the protein is detailed in our previous work (22). Briefly, expression strains were grown in LB + Kan (50 μg/mL) to OD600 0.6–0.8, induced with IPTG (0.2 mM) at 16 °C for 16 h, and harvested by centrifugation. Cells were resuspended in lysis buffer and lysed, and the supernatant was then collected. Recombinant FtsZ purification involved Ni^2+^-NTA affinity chromatography with elution using Tris-HCl and 300 mM imidazole, followed by Q-anion exchange using 0–1 M NaCl gradient, concentration with a 30-kDa ultrafiltration tube, and final purification by Superdex^TM^ 200 size-exclusion chromatography. Protein concentration was measured with the BCA assay, and purity was determined using SDS-PAGE. The purified protein was frozen in liquid nitrogen and kept at -80 °C.

### 3D Structure Prediction of SmFtsZ

The 3D structure of SmFtsZ was predicted with the advanced protein modeling tool AlphaFold 3 (37). The SmFtsZ truncation spanning residues 1–319 was selected as the prediction model because of the very low per-residue local distance difference test (LDDT) confidence scores of the C-terminal peptide. The DNA primary sequence and GTP were directly input into AlphaFold 3, which produced five structural models in its default mode. Of these, the model with the highest prediction confidence score referred to as “_model_0,” was chosen for presentation. The cartoon representation of this structure, as visualized on the web server, was saved and included in the current study.

The AlphaFold Server is accessible at https://alphafoldserver.com/.

### Crystallization, Data Collection, and Structure Determination

Purified SmFtsZ truncation 1–319 (∼14 mg/mL) was incubated with 2 mM GTP for 15 min. Crystallization trails were performed at 20 ℃ by the hanging-drop vapor-diffusion method. The crystallization condition used for determining the structure of GTP-bound SmFtsZ contained 0.1 M MES (pH 6.7), 8% (w/v) PEG10,000, and 0.1 M potassium sodium tartrate (Seignette salt). The SmFtsZ-GTP complex crystals were generated within four days and rapidly frozen in liquid nitrogen at 77 K using 25% (w/v) ethylene glycol as a cryoprotectant. The X-ray diffraction data for a single crystal of the SmFtsZ-GTP complex were collected at Shanghai Synchrotron Radiation Facility (SSRF) on beamline BL02U1 and processed using the XDS package. Molecular replacement searches were conducted using Phaser MR with the structure of FtsZ (PDB: 2RHL) as the starting model. Phenix.refine and WinCoot facilitated the processes of model construction and refinement. MolProbity was used for structure validation; statistics for data processing and refinement are shown in Table 1. Structure visualization was performed with PyMOL 3.1.3.

### Small-angle X-ray scattering (SAXS)

Data collection parameters and analysis software were consistent with previously described methods (38). Protein samples (final concentration, 4 mg/mL), were purified by gel filtration chromatography in buffers with varying pH values: (1) 50 mM Tris–HCl pH 7.4, 250 mM KCl, 5 mM MgCl_2_, 3% glycerol and, 2mM DTT; (2) 50 mM MES pH 6.0, 250 mM KCl, 5 mM MgCl_2_, 3% glycerol and, 2mM DTT. A PILATUS 100 k detector (Dectris) was used to record scattered X-ray photons at BL19U2. Experimental setups were configured to achieve scattering q-values of 0.007 < *q* < 0.450 Å−1, where *q* = (4π/λ)sin(θ), and 2θ is the scattering angle. Twenty two-dimensional (2D) images were recorded for each sample solution or corresponding buffer using a flow cell, with an exposure time of 1 s. Protein scattering profiles were determined by removing the background buffer contribution from the sample buffer profile with PRIMUS3.2 (39) according to established protocols. The radius of gyration (*R*_g_) and maximum dimension of the molecule (*D*_max_) were estimated from the scattering profiles with a broader q range of 0.007–0.30 Å^−1^ using the indirect Fourier transform method implemented in the program GNOM. The volume of correlation (V_c_) was determined using the Scatter program, and the molecular weights of solutes were calculated using the R_g_/V_c_ power law (40).

### Development of the FtsZ Amino Acid Mutation Strain

In this study, we constructed a fusion strain of the FtsZ 68 arginine mutant protein using the IFDC2 dual-screening system. The IFDC2 DNA fragment contains a *ldh* promoter, an erythromycin resistance gene, and a modified *phes\** gene. The transcription and translation product of the *phes\** gene acetylates p-chlorophenyl alanine (*p-Cl-Phe*), which exerts a toxic effect on bacterial cells, leading to their death. Consequently, when IFDC2 is integrated into the bacterial genome, positive clones can be selected based on erythromycin resistance. After removing IFDC2, positive clones are identified using medium supplemented with *p-Cl-Phe*. To counteract the potential lethality caused by the loss of the FtsZ protein, we introduced the pZX10-FtsZ plasmid, enabling FtsZ expression upon xylose induction.

We identified the *ftsZ* gene and approximately 2000 base pairs of its flanking sequences in the *S. mutans* UA159 genome from NCBI. Primers were designed based on this sequence to amplify 800–1000 base pair DNA fragments upstream and downstream of *ftsZ*. The amplified upstream fragments, *ftsZ-R68A*, mNeogreen fluorescent sequences, and downstream fragments were combined using fusion PCR. The resulting fusion product was then used to transform the FtsZ knockout strain of *S. mutans* using the CSP peptide to induce bacterial cell transformation. The strain expressing the mutant FtsZ protein at the native *F*tsZ locus was selected via spectinomycin-resistant plates.

### FtsZ Polymerization and GTPase Assay

FtsZ and FtsZ-mutations (10 μM) were polymerized in a buffer containing 5 mM Mg^2+^, 150 mM KCl, and 2 mM GTP at specific pH levels (pH 7.4 with Tris, pH 6.0 with MES, and pH 5.0 with sodium citrate) for 15 min at room temperature. After polymerization, a 1% uranyl acetate solution was used to stain the samples, which were then analyzed using an electron microscope (JEOL JEM-F200, JEOL Ltd., Tokyo, Japan).

Simultaneously, GTPase activity was assessed by quantifying the release of inorganic phosphate, which correlates with GTP hydrolysis. An assay based on malachite green was used to detect levels of inorganic phosphate. FtsZ and its mutants were prepared in a polymerization buffer containing 5 mM MgCl_2_, 150 mM KCl, and 50 mM of various pH buffer solutions within 96-well plates. The reaction was initiated by adding 1 mM GTP and incubating for 5 min. Phosphate release was then detected using the chromogenic agent MAK307-1KT (Sigma, USA), and absorbance was measured at 620 nm with a full-wavelength microplate reader.

### Imaging *S. mutans* FtsZ_mNeongreen dynamics

In this study, FtsZ_mNeongreen fluorescent protein was imaged using an N-STORM microscope, with illumination at an angle near the total internal reflection fluorescence (TIRF) to selectively activate fluorescent proteins at the base of the bacteria. As the fluorescent proteins move through the imaging area, they create a continuous dynamic image reflecting changes in fluorescence intensity. Fluorescent proteins were excited with a 488 nm laser, using an exposure time of 0.166 s and an interval of 0.83 s, over a total imaging period of 10 min.

We used Fiji (version 2.1.0/1.53c) to analyze the “treadmilling activity” of SmFtsZ and its mutations. The built-in plugin Descriptor-based series registration (2d/3d+t) in Fiji was used to correct for image drift, and a 3×3 pixel area (∼320 nm) within the Z-ring was selected. The plot of the Z-axis profile function in Fiji was then used to analyze variations in fluorescence intensity within this area over time. A straight line was drawn along the long axis of the interior of the Z ring, and the kymograph analysis tool (https://www.embl.de/eamnet/html/body_kymograph.html) was applied to assess changes in fluorescence intensity along this line. The macro “read velocities from tsp,” provided in the multiple kymograph plugin manual, was used to analyze the kymograph images and calculate the movement velocity of FtsZ.

### Snapshotting images of FtsZ

*S. mutans* FtsZ_mNeongreen was imaged using the N-STORM system (Nikon, Tokyo, Japan), which includes a 100 ×oil immersion TIRF objective lens (Nikon Plan Apo, 1.49 NA) (32), the Andor-897 EMCCD imaging platform (Andor, Belfast, Northern Ireland), lasers emitting at 405, 488, 561, and 647 nm, and optics providing 1.5 × magnification. Image analysis was performed using ImageJ software (NIH, Bethesda, MD, USA). Image stacks in .nd2 format acquired via Nis-Elements AR software were converted into .tiff files, and Z-projections were generated in ImageJ. The images underwent further analysis to extract data on cell outlines and fluorescence signals. Bright-field images were used to manually delineate cell outlines. Custom MATLAB scripts (https://github.com/interestinghua/MapZ-and-FtsZ-rings/tree/master) were used to analyze the integrated fluorescence signal profile and locate the maximum intensity, indicating the site of FtsZ ring localization. Z-ring thickness was then manually measured in ImageJ.

### Transmission Electron Microscope Analysis

*S. mutans* UA159 and the SmFtsZ-R68A mutation strain were cultured in CDY medium at pH 7.4 and 5.0 until the absorbance at 600 nm (OD600) approached 0.4. The cells were harvested by centrifugation at 3,220 ×g for 15 min, rinsed once with PBS, and preserved in 2.5% glutaraldehyde/2% paraformaldehyde at room temperature for 1 h. Following a PBS wash, the cells were treated with 1% osmium tetroxide for 1 h and then rinsed in water. The samples were stained en bloc using 1% uranyl acetate for 1 h. After washing and dehydration through a graded ethanol series, the cells were embedded in Eponate 12 resin. Ultrathin sections (∼90 nm-thick) were prepared, stained with uranyl acetate and lead citrate, and observed using a JEOL 1400 EX transmission electron microscope.

### Growth Curve Analysis

The growth of test strains was monitored using a SpectraMax 190 Microplate Reader (Molecular Devices, Sunnyvale, CA, USA) and analyzed with corresponding software. Overnight cultures were diluted 1:100 in CDY medium with an uninoculated control. Growth was tracked by recording OD600 at 30-minute intervals over 24 h at 37 °C, with vigorous shaking for 5 sec prior to each measurement. Growth curves were generated by plotting OD600 against time.

### Biofilm Analysis and Structural Imaging

*Biofilms of S. mutans* were cultivated in CDY medium (pH 7.4). The cells were labeled using a combination of SYTO 9, which fluoresces green for live cells, and propidium iodide, which fluoresces red for dead cells. Biofilm visualization was performed using a confocal laser scanning microscope (TCS-SP8 and MICA, Leica) equipped with a 25× water immersion objective lens. Imaging gates were set to 480/500 nm for SYTO 9 and 490/635 nm for propidium iodide. Biofilms were scanned at five random positions, and optical sections were used to create 3D reconstructions of the biofilm structure. Representative image stacks from three independent experiments were selected for analysis. The survival rate of live cells was quantified using NIS Elements imaging software (NIS-Elements, Tokyo, Japan).

### Dental caries animal model

All animal experiments were conducted with approval from the Ethics Committee of Peking University, China (PUIRB-LA2024229). The dental caries animal model was established using male SPF-SD rats weaned at 21 days of age. During the initial phase of model establishment (days 1–4), the rats were administered streptomycin in their drinking water. From days 5 to 10, the oral cavities of the rats were inoculated with both the WT and mutant strains of *Streptococcus mutans*. On day 10, saliva samples from the rats were diluted and spread onto Brain Heart Infusion (BHI) agar to assess bacterial colonization. The agar plates were then kept in a 5% CO_2_ environment at 37 °C for 48 h to confirm successful colonization.

The rats were then maintained on a high-sugar diet for 40 days. On day 50, they were humanely euthanized using CO_2_ gas, and their skull specimens were harvested. The collected specimens were subsequently subjected to micron-scale computed tomography (micro-CT) imaging.

### Keyes’ Scoring Method for Evaluating Caries

Following the micro-CT examination, the samples were prepared for analysis using the revised Keyes’ scoring technique. Each specimen was treated with a 0.4% ammonium purpurate solution and allowed to stain for 24 h. The stained samples were then bisected along the mesiodistal orientation using a precision cutter with a thickness setting of 0.1 mm. A high-definition stereomicroscope (Olympus SZ61, Olympus Corporation, Tokyo, Japan) was used to assess the extent of the cavity in these specimens. Under microscopic observation, carious lesions appeared red, while unaffected, sound, and hard dental tissues remained unstained. Cavity decay was assessed by linearly estimating the depth of the cavity as if spread flat. The total linear extent was divided into three equal parts (see Fig. 4*D*). The severity of the lesion was quantified based on the number of these units affected, ranging from zero to three. Both sides of each bisected jaw section were examined during the assessment (41).

### RNA Sequencing

*S. mutans* UA159 and its mutant strains were grown to the exponential phase in CDY medium (pH 5.0) at 37 °C. Bacterial pellets were then collected for RNA extraction. Total RNA extraction was performed with the TRIzol reagent (Invitrogen, CA, USA) following the manufacturer’s instructions. RNA purity and concentration were measured using a NanoDrop 2000 spectrophotometer (Thermo Scientific, USA), and RNA integrity was evaluated with an Agilent 2100 Bioanalyzer (Agilent Technologies, Santa Clara, CA, USA). Samples meeting the required quantity, integrity, and purity standards were chosen for library construction. To remove ribosomal RNA, we used the TIANSeq rRNA Depletion Kit (TIANGEN, Beijing, China). Libraries were constructed using the VAHTS Universal V6 RNA-seq Library Prep Kit following the manufacturer’s directions. Transcriptome sequencing and analysis were conducted by OE Biotech Co., Ltd. (Shanghai, China).

### Molecular Dynamics Simulation

Based on the crystal structure of the SmFtsZ tetramer, we modeled the initial dimeric conformations of SmFtsZ with longitudinal SmFtsZ-dimer_long_ and lateral SmFtsZ-dimer_late_ interaction interfaces (Fig. S5). In the dimeric complex, one GTP molecule along with a Mg^2+^ ion was considered to bind with one SmFtsZ monomer at the dimeric interface. The longitudinal interaction interface is located where the protein molecules meet at the head and tail, whereas the lateral interaction surface is formed around the axis. We performed all-atom CpHMD simulations (23) of SmFtsZ-dimer_long_ and SmFtsZ-dimer_late_ in explicit solvent with a model for discrete protonation states at pH values 5.0, 6.0, and 7.0. During the simulations based on the pKa values, we defined the discrete protonation states of the titratable acidic residues aspartate (D), glutamate (E), and histidine (H) in each monomer of the SmFtsZ protein. The pKa values were considered as follows: D ∼ 4.0, E ∼ 4.4, and H ∼ 6.6. The D, E, and H residues of each SmFtsZ monomer within 20 Å of the dimeric interface were considered titratable in the dimeric complexes. The fully solvated systems of SmFtsZ-dimer_long_ and SmFtsZ-dimer_late_ using the tLEAP module of Amber Tools 18. The systems included an orthorhombic water box containing TIP3P water molecules and 89 Na^+^ and 50 Cl^−^ ions randomly placed to neutralize the system. A layer of water molecules at least 15 Å thick surrounded the solute in all directions. CpHMD simulations, lasting 500 ns, were conducted at pH values 5.0, 6.0, and 7.0 to characterize the binding interactions at the dimeric longitudinal and lateral interfaces under acidic (pH) conditions. We also introduced the R68A mutation into the SmFtsZ-dimer_late_ complex and performed simulations at pH 5.0 and 6.0.

CpHMD simulations of SmFtsZ-dimer_long_ and SmFtsZ-dimer_late_ were performed using the method developed by Swails et al.(23), using AMBER 18 and parametrized with AMBER ff14SB force field (42). Each solvated system was energy minimized for 20,000 steps using a combination of steepest descent and conjugate gradient algorithms under periodic boundary conditions. The minimized systems were then heated to the desired temperature (300K) over 100 ps under constant volume (NVT) conditions, with weak restraints on the solute atoms, using Langevin dynamics with a friction coefficient of 1 ps^−1^. The systems were then equilibrated for 500 ps under constant pressure (NPT) simulation at 1 atm to adjust to the appropriate volume.

The particle-mesh Ewald method with default parameters was used to model long-range electrostatic interactions accurately (43). After equilibration, we conducted 500 ns CpHMD simulations of each system at constant pressure and temperature using a 2 fs time step at pH values 5, 6, and 7. The CpHMD method involved propagating standard MDS in explicit solvent, with attempts to modify protonation states using generalized Born (GB) model for implicit solvent at regular intervals. The GB model by Onufriev et al.(44) (with igb set to 2 in the sander model) was used to assess changes in protonation states, which were tested every 200 fs during all constant pH simulations. If a change in protonation state was accepted, the solute was immobilized. MD simulations were subsequently conducted on the solvent and ions to equilibrate the solvent distribution around the new protonation states. Solvent relaxation dynamics were performed using a relaxation length (τ_rlx_) of 200 fs. Once relaxation had completed, the solute atom velocities were reverted to their values prior to relaxation, and standard dynamics resumed (23). All MD simulations, including the relaxation of the solvent, were performed using a 2 fs time step, and hydrogen-containing bonds were constrained with the SHAKE algorithm (45). Conformations in each MD run were saved every 1 ps from the trajectories for further analysis.

Equilibration of each system at its respective pH was ensured by monitoring the root-mean-square displacement (RMSD) of the SmFtsZ dimer structure relative to its initial energy minimized initial structure. The final 200 ns of equilibrated trajectories were used for subsequent analyses presented here. Trajectory analysis was performed using the CPPTRAJ module of AMBER 18.

## Supporting information

20250127 manus...t-final

## Data availability

The SmFtsZ structure was submitted to wwPDB (ID 9LFX). The RNA-Seq data are available in Sequence Read Archive (SRA) under BioSample accessions PRJNA1206306. (https://www.ncbi.nlm.nih.gov/sra/PRJNA1206306)

## Acknowledgments

This work was supported by the Beijing Natural Science Foundation: 7222220, Research Foundation of Peking University School and Hospital of Stomatology: PKUSS20230117. We express our gratitude to the Tsinghua University Branch of the China National Center for Protein Sciences Beijing and to Shilong Fan for providing facility support in conducting X-ray diffraction on the crystal samples. We thank the Core Facilities of Life Sciences and the National Center for Protein Sciences at Peking University in Beijing, China, for their assistance with TEM imaging, and we are grateful to Rui Jiao and Yiqun Liu for their assistance in image acquisition. We appreciate the help of Dr. Jiao Liu from the Center of Medical and Health Analysis, Peking University Health Science Center, in confocal microscopic imaging. Computational resources were supported by the Shenzhen Bay Laboratory Supercomputing Centre.

