## Supplementary material for "Particular Amino Acid on Lateral Interface Maintains FtsZ Function Under Acid Stress in *Streptococcus mutans*": 20250127 manus...t-final

**This PDF file includes:**

Supporting text
Figures S1 to S15
Tables S1 to S5

**Supporting Information Text**

**Molecular Dynamics simulation studies on interactions at the binding interface of SmFtsZ in acidic pHs environments**

**Binding interface & stability of secondary structural elements of SmFtsZ in dimeric complex.** From our all-atom MD simulation studies SmFtsZ chains are found to maintain the dimeric assembly at different pHs through interactions among the interface residues as depicted in the residue contact maps (Fig. S13). We found that residues 134-150 and 165-190 of chain-A, which includes H5 and H6 helices mainly form the interface contacts with the residues 200-222 and 258-300 of chain-B that contains H8 and H10 helices, S8 and S9 sheets and T7 loop in SmFtsZ-dimer$_{long}$ case (Fig. S5, S13*A-C*). On the other side in SmFtsZ-dimer$_{late}$ complex, residues 40-96 contain H2 and H3 helices, S2 and S3 sheets and T3 loop in the N-terminal domain of chain-A form the interface contacts with the resides 224-314 include H9 and H10 helices in the C-terminal domain of chain-B (Fig. S5, S13*D-F*). Variation of secondary structural elements of SmFtsZ chains in SmFtsZ-dimer$_{long}$ and SmFtsZ-dimer$_{late}$ complexes over the simulation time (Fig. S14, S15) does not show any major change in folded conformation of protein chains at acidic pH environment. However, N-terminal H1 (21-34) and H2 (48-53) helices in GTP-binding pocket of chain-A, H8 (212-219) helix of chain-B, and H5 (142-158) and H6 (167-173) helices of chain-A and chain-B show differential stability at pH 5.0, 6.0 and 7.0 in SmFtsZ-dimer$_{long}$ complex (Fig. S14). Similarly, pH dependent different stability of H5, H6 and H7 (180-203) helices of both the protein chains are found in case of SmFtsZ-dimer$_{late}$ (Fig. S15).

**Interface area and binding interactions in SmFtsZ dimeric complex.** We analyzed the total buried surface area (BSA) and BSA of the individual protein chains, which measures the size of the interface in the dimeric complexes in different pH environment. In case of SmFtsZ-dimer$_{long}$, average total BSA is comparatively higher with greater contribution from chain-A at pH 6 (Fig. S9*A*). Average non-bonded interaction energy (Eint) is also comparatively high at pH 6 (Fig. S9*B*), where electrostatic interactions mainly favor the dimeric association. At the same time binding interaction of GTP with chain-A is more favored at pH 6 (Fig. S9*C*). As compared to the physiological pH, Eint between the protein chains in SmFtsZ-dimer$_{late}$ complex is higher at acidic pHs (Fig. S9*D*), and total BSA and BSA of the individual protein chains attain higher value at pH 5 (Fig. S9*E*). The average number of potential hydrogen bonds (NH) between the protein chains show comparatively higher value at pH 6 for SmFtsZ-dimer$_{long}$ and at pH 5 and 6 for SmFtsZ-dimer$_{late}$ (Fig. S9*F-G*).

**At different pH environment interactions among the key interface residues in (SmFtsZ-dimer)$_{long}$ complex.** We studied the key interactions among the interface residues in SmFtsZ-dimer$_{long}$ complex simulated at different pHs. Fig. S11*A-C* show the interface residues in SmFtsZ-dimer$_{long}$, which form favorable hydrogen bonds between the protein chains and stabilize the GTP in the binding pocket of chain-A at pH 5.0, 6.0 and 7.0. Arg144 of chain A forms salt-bridge with Asp214 of chain-B both at pH 7.0 and 6.0. At pH 7 backbone atoms of F139 and K143 of chain A form favorable hydrogen bonds with the L295 and N292 of chain-B, respectively (Fig. S11*A*). Besides, Asn285, Leu180 and Phe139 of chain-A form potential hydrogen bonded interactions with Lys150, Met271 and Ser280 of chain-B, respectively at pH-6 (Fig. S11*B*). Ile293 of chain-B also forms hydrogen bonded interactions with Ser142 and Lys143 of chain-A at pH 5.0 and 6.0. Salt-bridge interaction of Arg144-Asp214 is disrupted at pH 5.0 due to protonation of side chain of Asp214. However, protonation of side chain of Asp211 of chain-B and Glu140 of chain-A assists to form the hydrogen bonded contact with the phosphate group of GTP and helps to stabilize at acidic pHs (Fig. S11*B-C*).

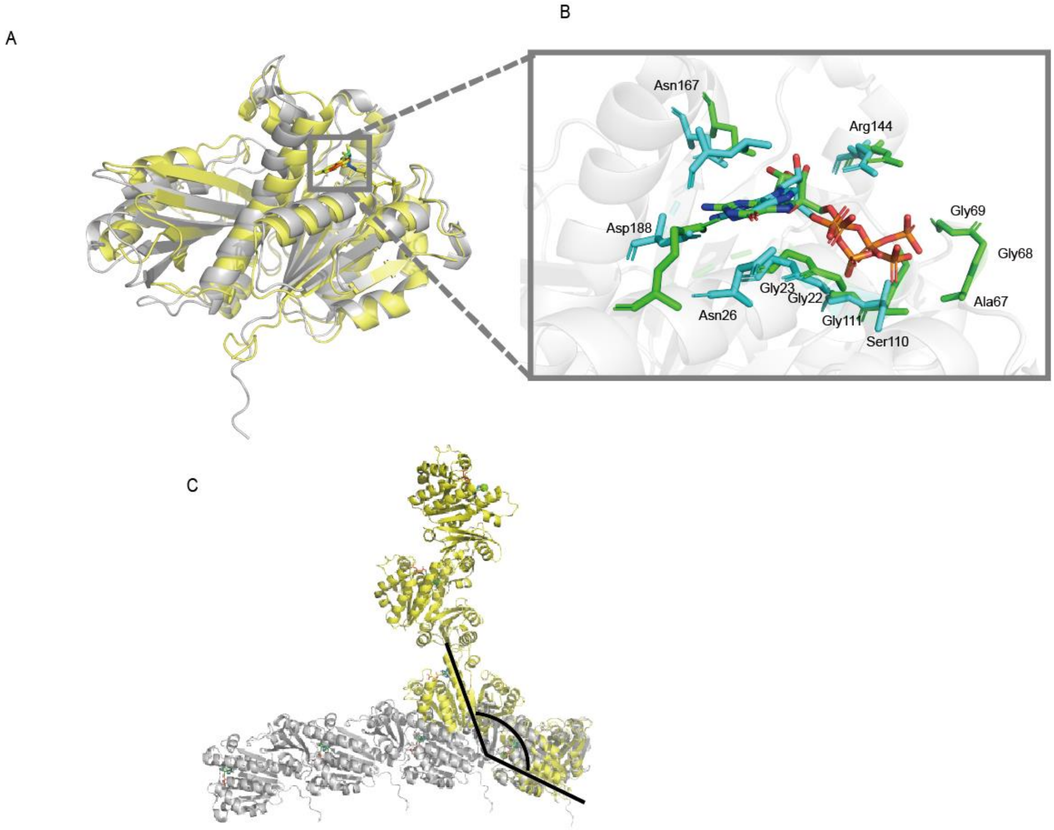


**Fig. S1.** (*A*) Superposed monomer of SmFtsZ solved from crystal structure and predicted by AlphaFold3. (*B*) GTP binding sites were shown. (*C*) Superposed tetramers of SmFtsZ solved from crystal structure and predicted by AlphaFold3.

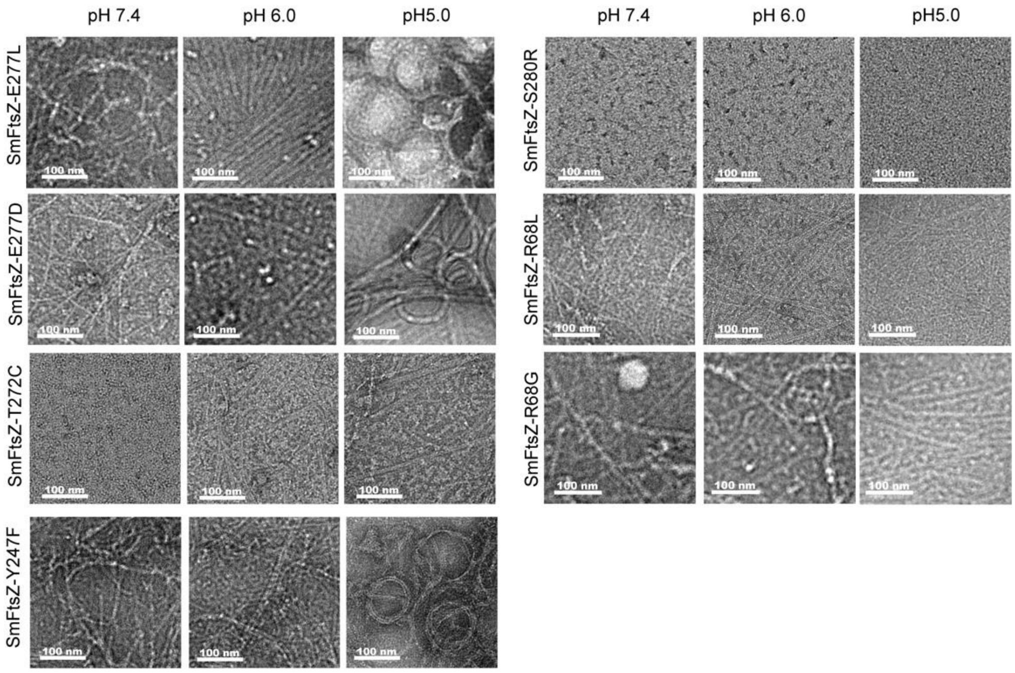


**Fig. S2.** Effect of pH on the polymerization of SmFtsZ variants.

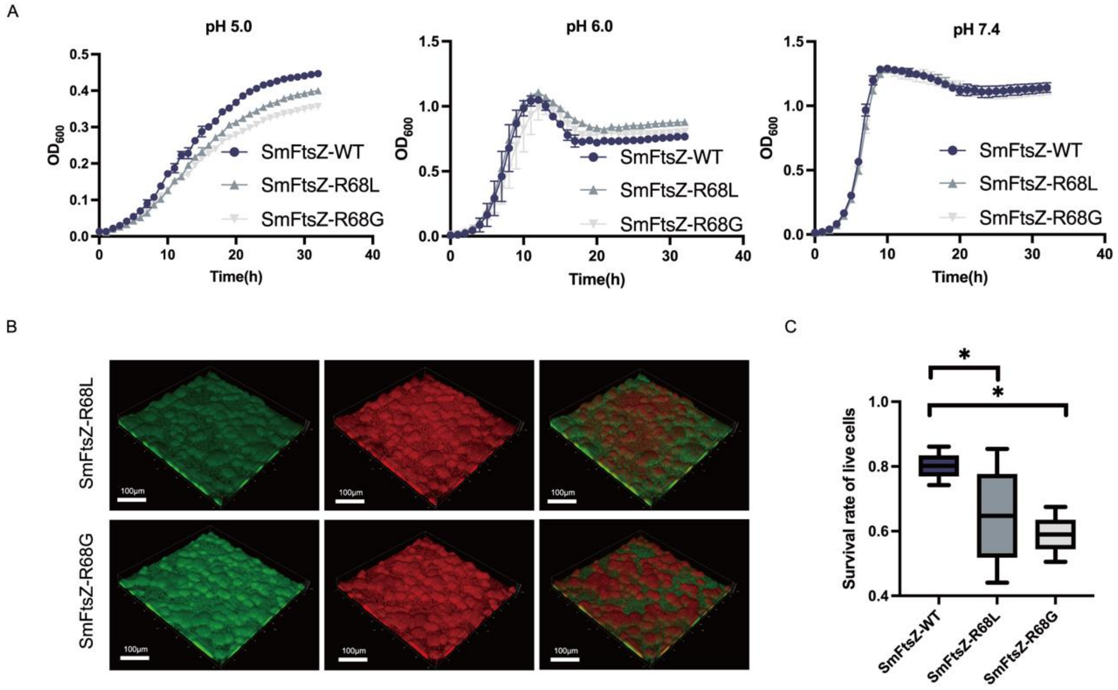


**Fig. S3.** Effects of an acidic environment on *S. mutans* UA159 strain and SmFtsZ-R68L/G strains. (*A*) Growth curves of *S. mutans* UA159 strain and SmFtsZ-R68L/G strains under different pH levels. Data were obtained from three independent experiments. (*B*) Representative CLSM images of biofilm of *S. mutans* UA159 strain and SmFtsZ-R68 L/G strains. Lived cells (green) were stained with SYTO 9 stain, and dead cells (red) were stained with propidium iodide. Images were examined at 25× objective magnification. Scale bar: 100 µm. (*C*) Survival rate of live cells of biofilm analysis using Leica imaging software. Data represent the means of three independent experiments. '*' indicates $p < 0.05$ (compared with SmFtsZ-WT respectively with t-test).

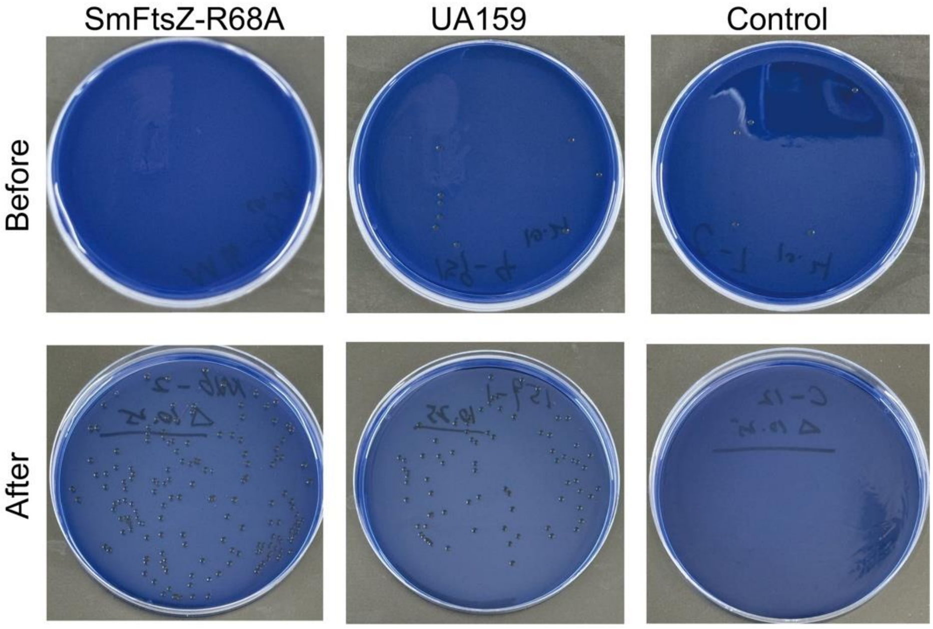


**Fig. S4.** Colonies of S. mutans UA159 and SmFtsZ-R68A strain were successfully detected after oral inoculation.

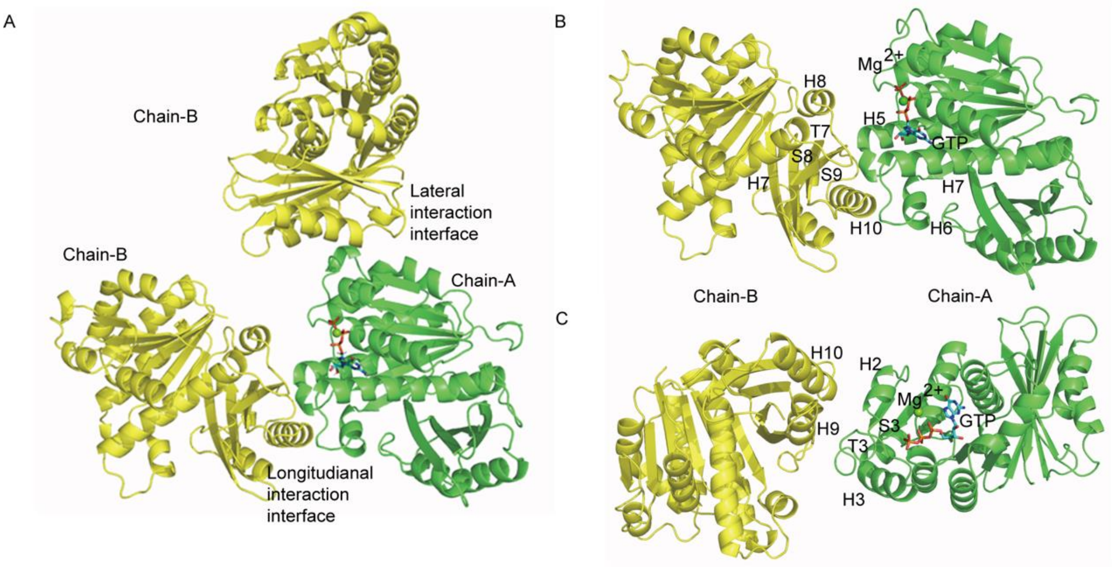


**Fig. S5.** Model structure of SmFtsZ dimer with longitudinal SmFtsZ-dimer$_{long}$ and lateral SmFtsZ-dimer$_{late}$ interaction interfaces(*A*), and structure of (*B*) SmFtsZ-dimer$_{long}$ and (*C*) SmFtsZ-dimer$_{late}$ for molecular dynamics simulations studies.

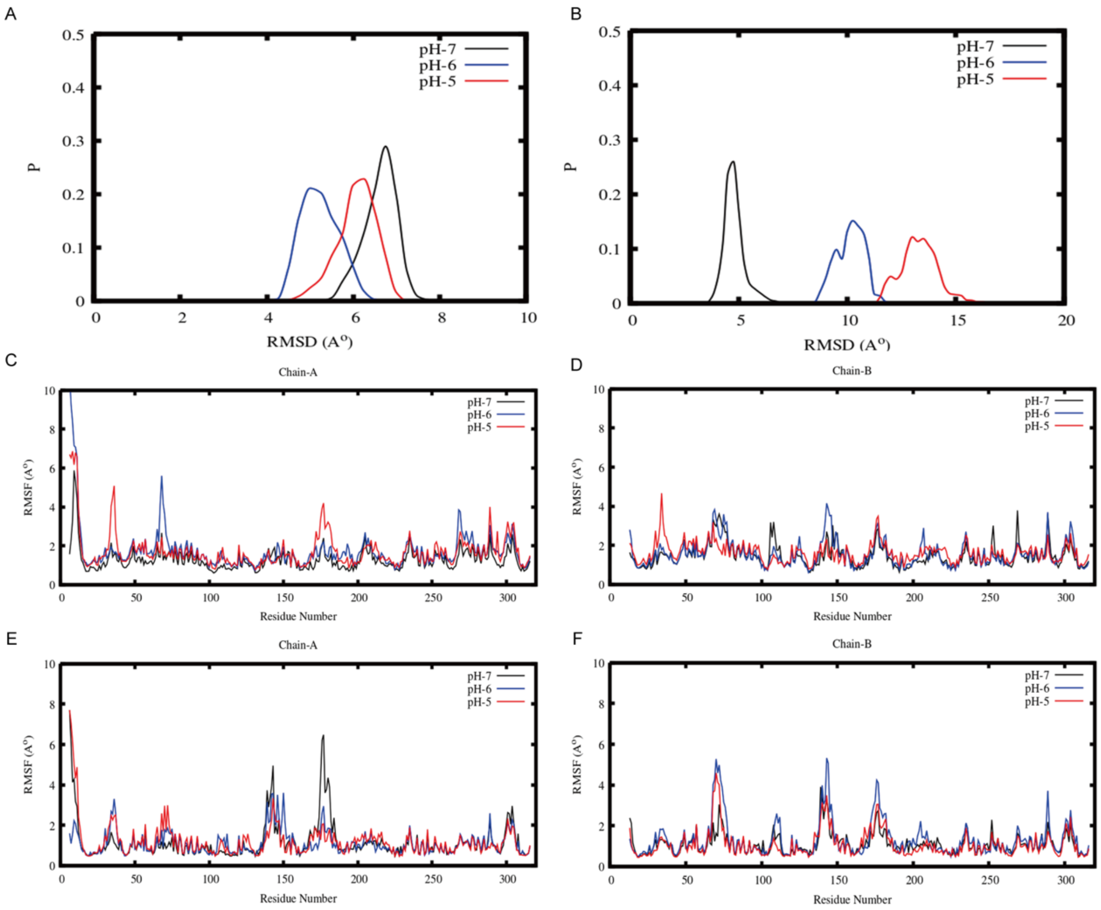


**Fig. S6.** Distribution of the root mean square deviations (RMSD) of the (*A*) SmFtsZ-dimer$_{long}$ and (*B*) SmFtsZ-dimer$_{late}$ structures with respect to the initial structure, and root mean square fluctuations of the residues in the individual protein chains in (*C*-*D*) SmFtsZ-dimer$_{long}$ and (*E*-*F*) SmFtsZ-dimer$_{late}$ complexes over the equilibrated MD trajectories at different pHs.

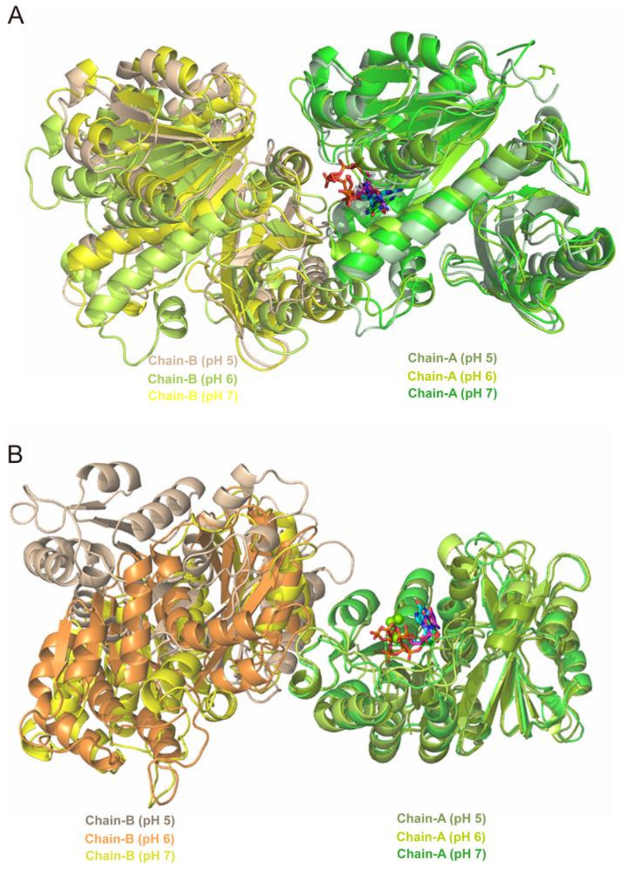


**Fig. S7.** Superposed simulated average structure of (*A*) SmFtsZ-dimer$_{long}$ and (*B*) SmFtsZ-dimer$_{late}$ at pH 7.0, 6.0 and 5.0.

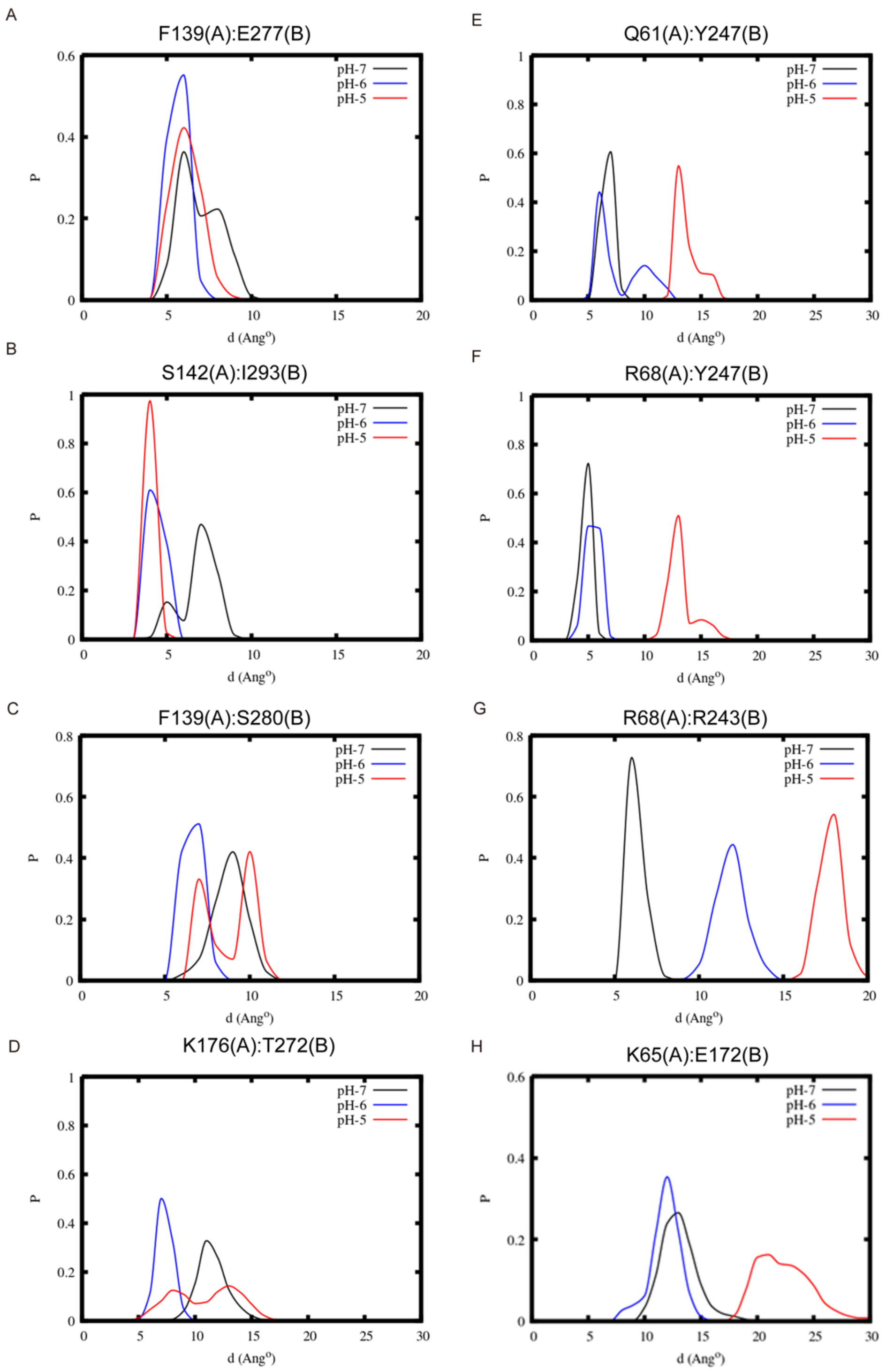


**Fig. S8.** Distribution of the distances of representative residues pairs of protein chains in case of (*A-D*) SmFtsZ-dimer$_{long}$ and (*E-H*) SmFtsZ-dimer$_{late}$ at different pHs, which primarily form the contacts in the dimeric interface, as found in the initial crystal structure.

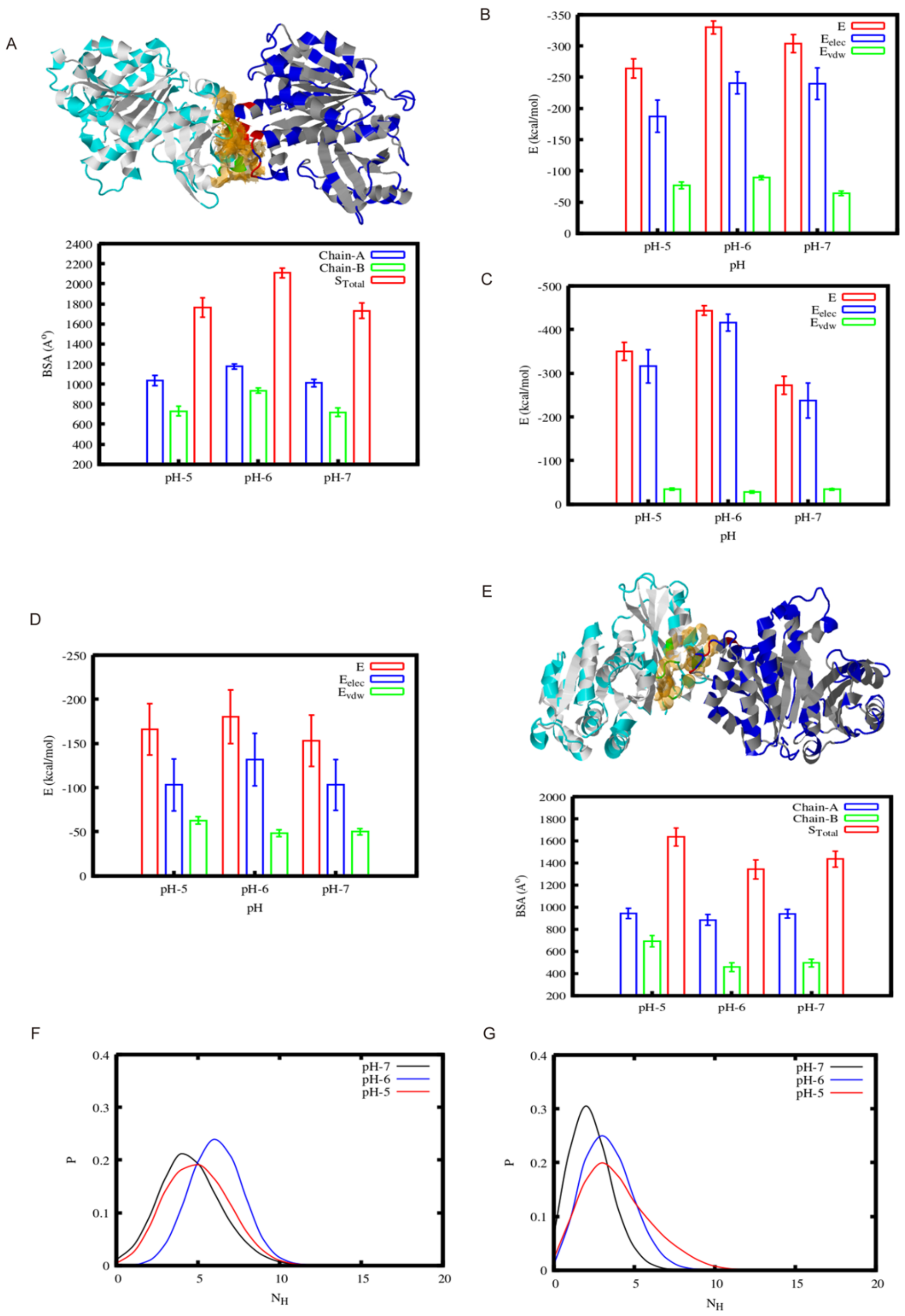


**Fig. S9.** In case of SmFtsZ-dimer$_{long}$ (*A*) total buried surface area (BSA) and BSA of the individual protein chains, and non-bonded interactions energy between the (*B*) protein chains and (*C*) chain-A with GTP at different pHs. (*D*) non-bonded interactions energy between the protein chains and (*E*) total BSA and BSA of the individual protein chains in SmFtsZ-dimer$_{late}$ complex at different pHs. Distribution of the number of favorable hydrogen bonds between the protein chains over the equilibrated simulated trajectories in case of (*F*) SmFtsZ-dimer$_{long}$ and (*G*) SmFtsZ-dimer$_{late}$ at different pHs.

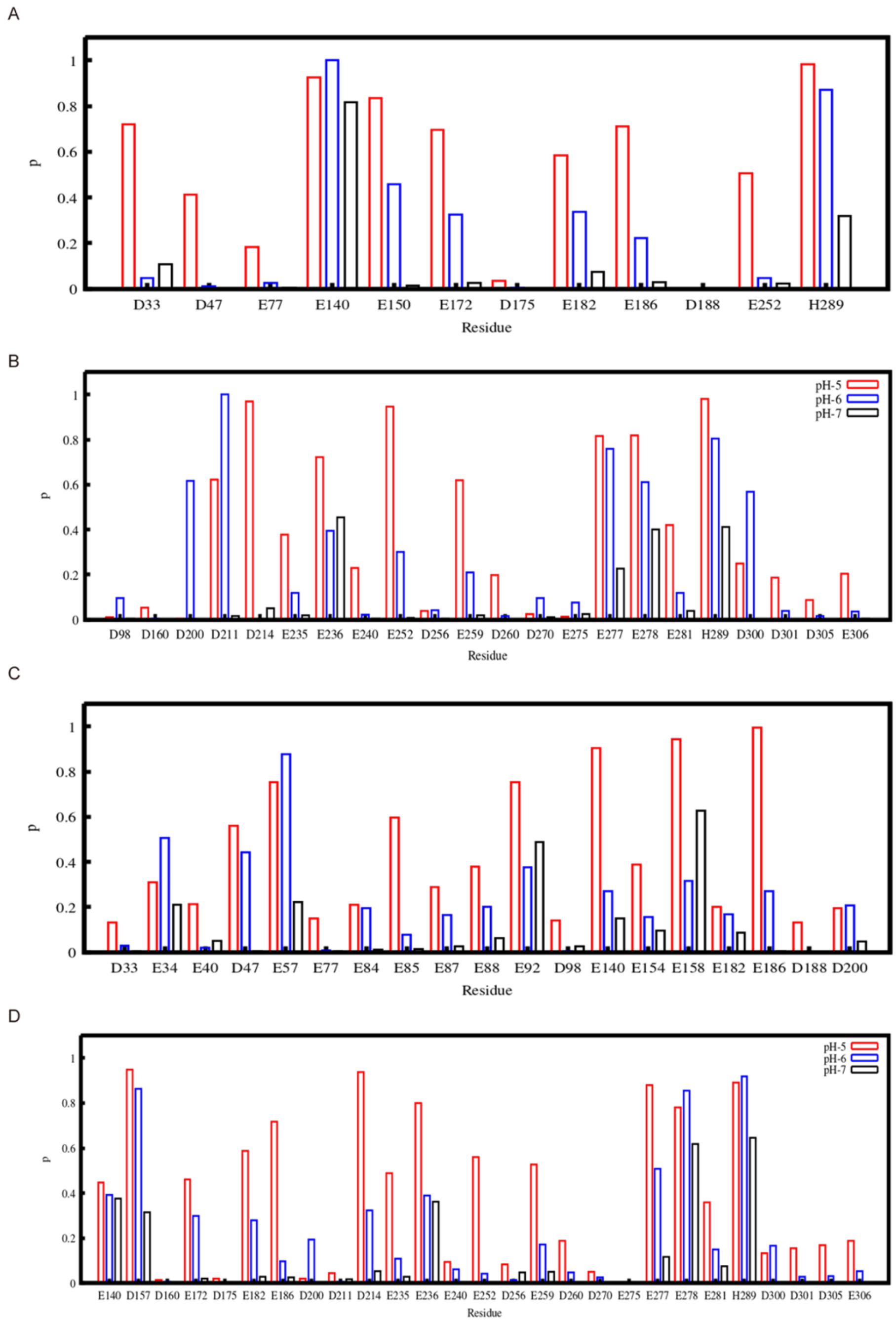


**Fig. S10.** Protonation probability of the titrated Asp, Glu and His residues of chain-A and chain B at pH 7 (black), 6 (blue) and 5 (red) over the equilibrated simulated trajectories in case of (*A*-*B*) SmFtsZ-dimer$_{long}$ and (*C*-*D*) SmFtsZ-dimer$_{late}$.

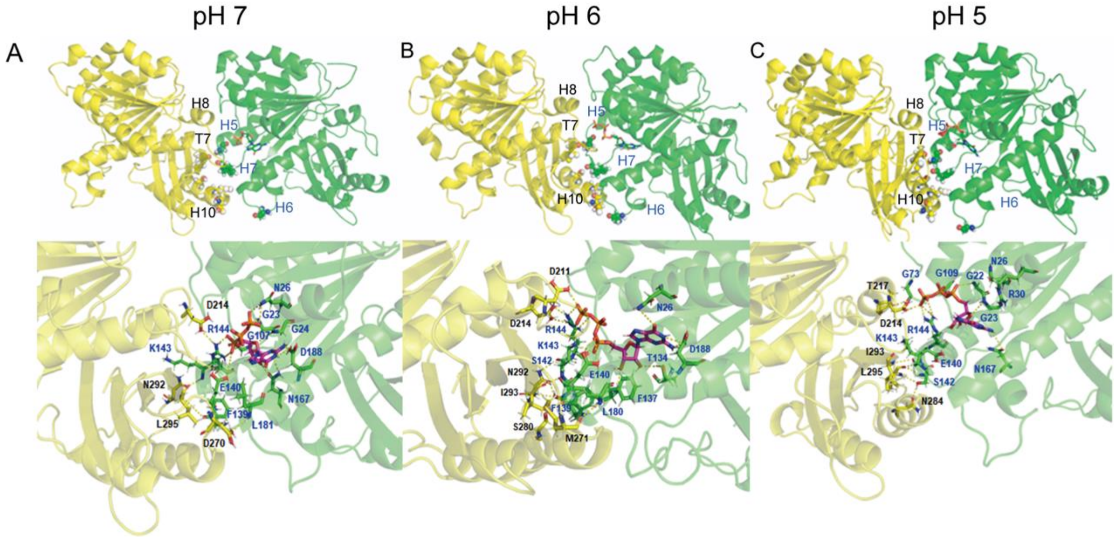


**Fig. S11.** Dimeric binding interface and interface residues, which form potential hydrogen bonds between the protein chains in (*A*-*C*) SmFtsZ-dimer$_{long}$, complexes at pH 7.0, 6.0 and 5.0, respectively.

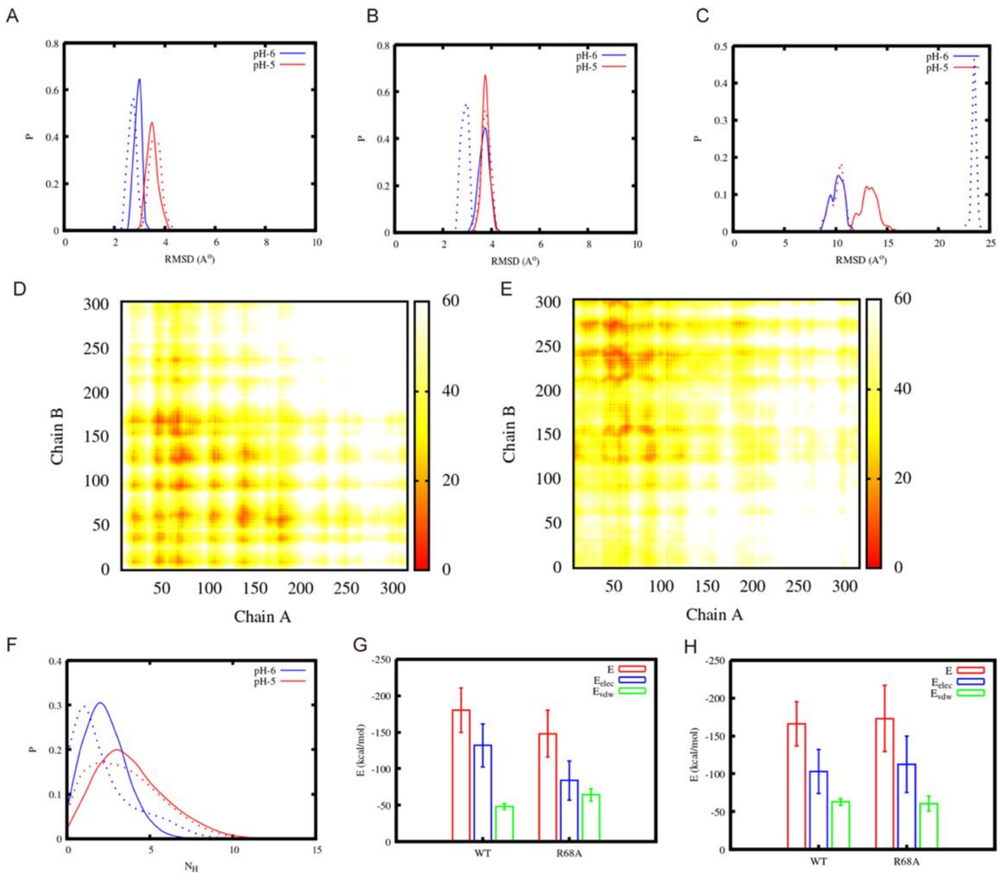


**Fig. S12.** Distribution of the root mean square deviations (RMSD) of the (*A*-*B*) individual protein chains and (*C*) dimeric lateral complex for wild type (solid line) and R68A mutated (dotted line) case with respect to the initial structure over the equilibrated MD trajectories at pH 5.0 and 6.0. (*D*) and (*E*) residues contacts map between the protein chains in dimeric lateral complex for R68A mutated case over the equilibrated MD trajectories at pH 6.0 and 5.0, respectively. The color bar represents the pair distance between the residues of protein chains. (*F*) Distribution of the number of favorable hydrogen bonds between the protein chains in lateral dimeric complex over the equilibrated simulated trajectories at pH 6 and 5 for SmFtsZ-WT (solid line) and SmFtsZ-R68A (dotted line) case. (*G*-*H*) Non-bonded interaction energy between the protein chains in lateral dimeric complex for SmFtsZ-WT and SmFtsZ-R68A case at pH 6.0 and 5.0, respectively.

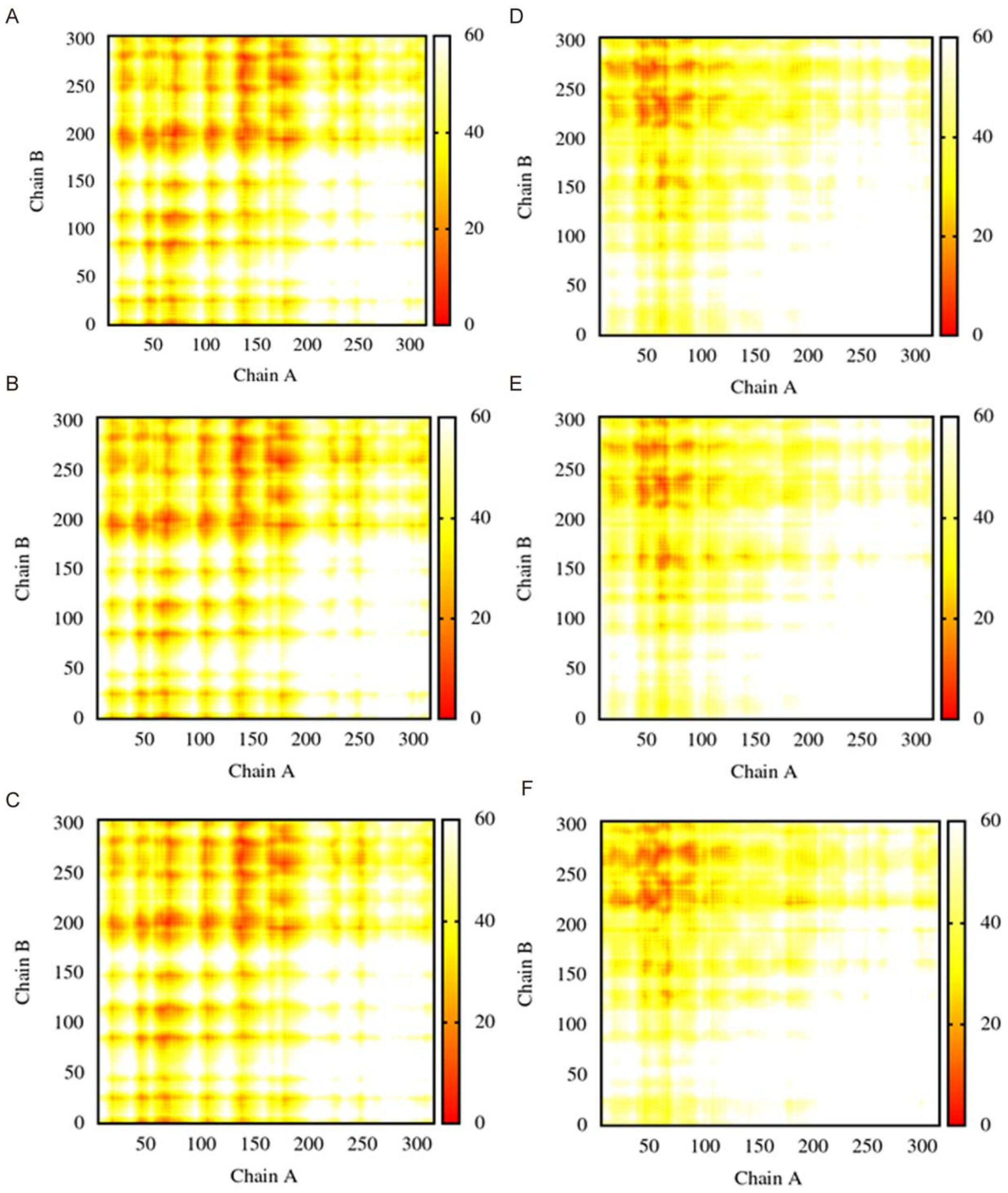


**Fig. S13.** Residues contacts map between the protein chains in (*A*-*C*) SmFtsZ-dimer$_{long}$ and (*D*-*F*) SmFtsZ-dimer$_{late}$ complexes over the equilibrated MD trajectories at pH 7.0, 6.0 and 5.0, respectively. The color bar represents the pair distance between the residues of protein chains.

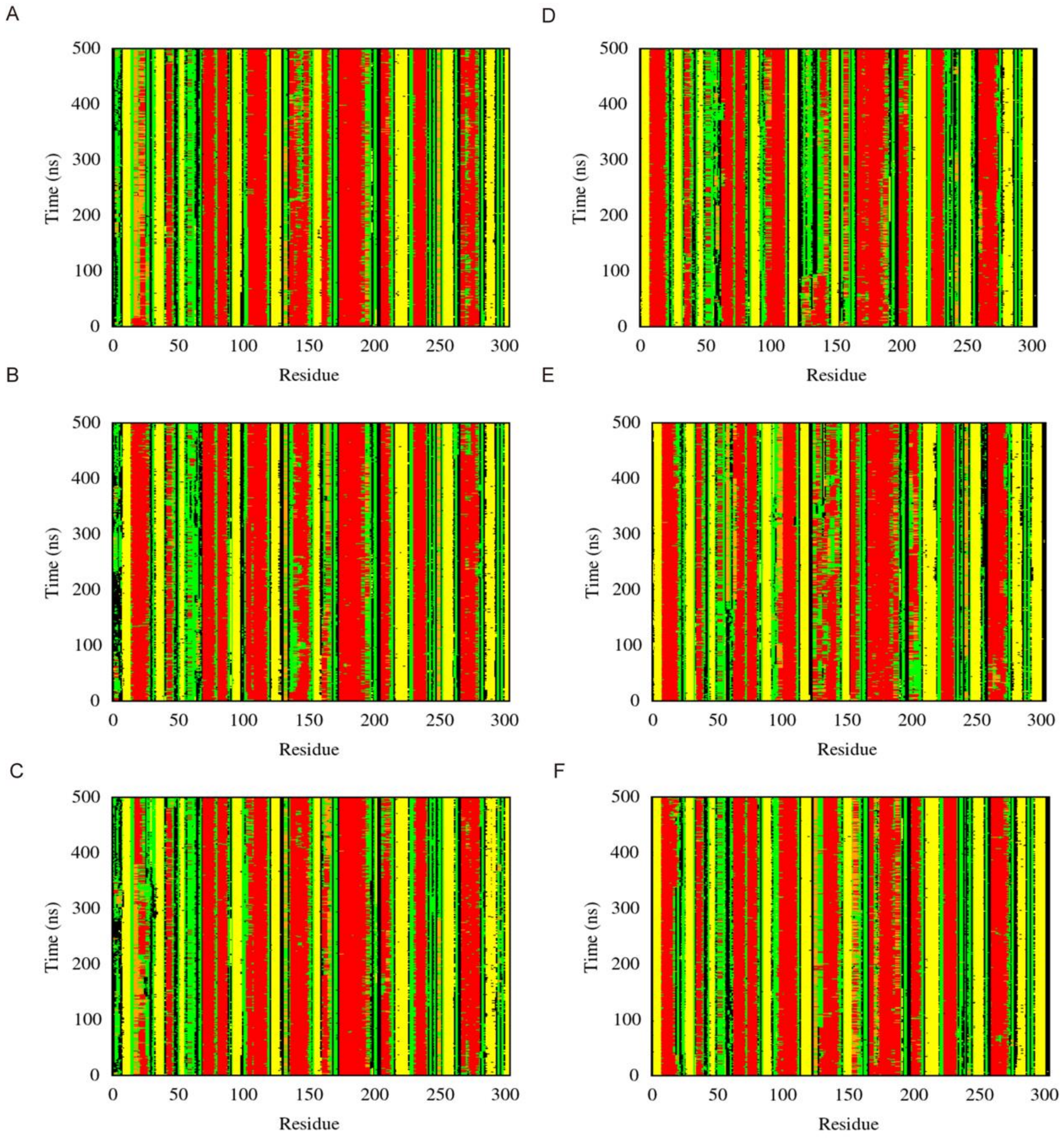


**Fig. S14.** Secondary structural preference of (*A*-*C*) chain-A and (*D*-*F*) chain-B at pH 7.0, 6.0 and 5.0, respectively over the simulation time in SmFtsZ-dimer$_{long}$ complex. (Secondary structural components are represented as α-helix: brown, 3-10 Helix/pi-Helix: orange, β-sheet: yellow, Turn/Bend: green).

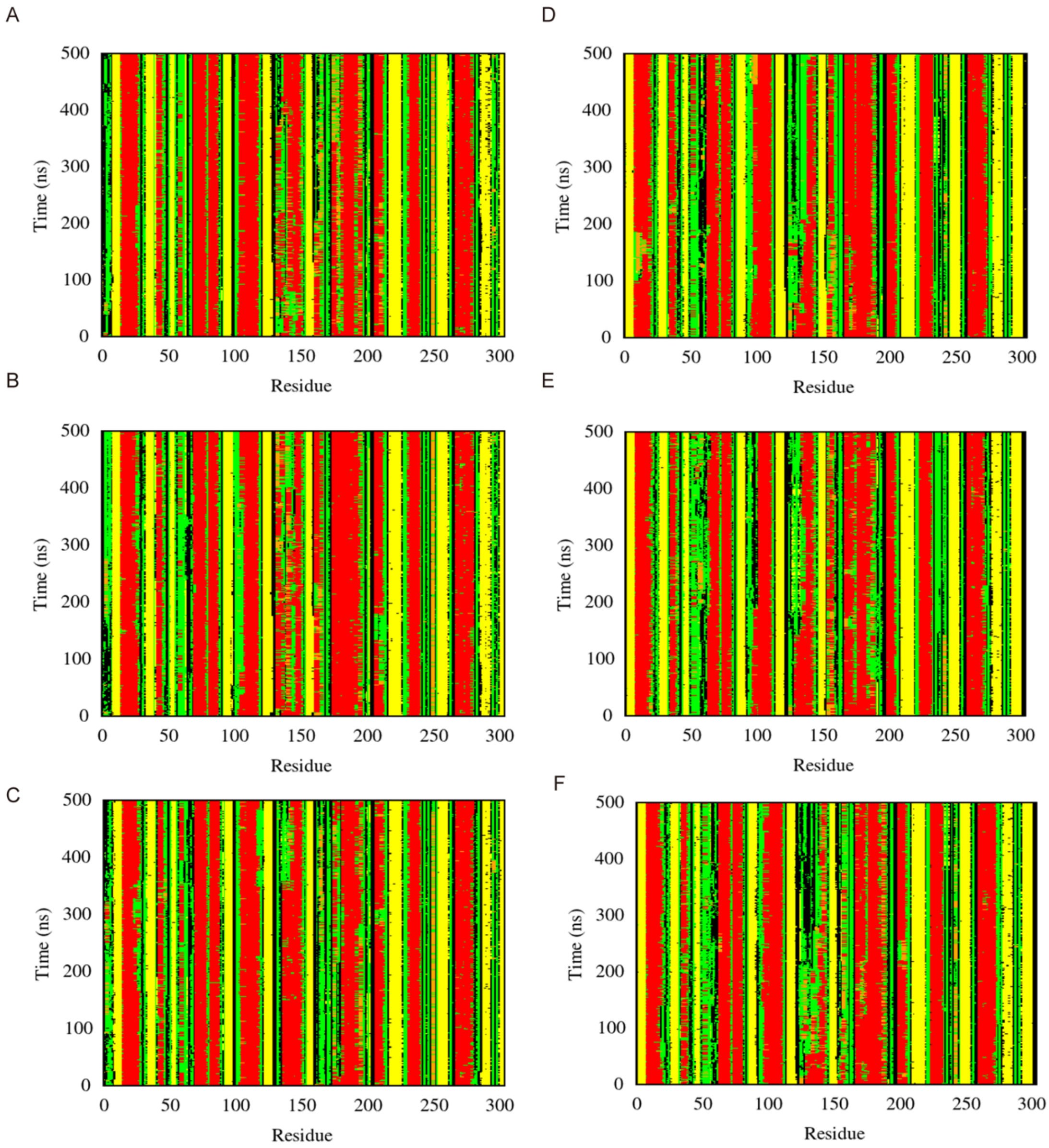


**Fig. S15.** Secondary structural preference of (*A-C*) chain-A and (*D-F*) chain-B at pH 7.0, 6.0 and 5.0, respectively over the simulation time in SmFtsZ-dimer$_{late}$ complex. (Secondary structural components are represented as α-helix: brown, 3-10 Helix/pi-Helix: orange, β-sheet: yellow, Turn/Bend: green).

**Table S1.** Data collection and refinement statistics

| Data collection and refinement statistics | |
|---|---|
| **Data collection** | |
| Space Group | $P4_32_12$ |
| Cell dimensions | |
| a, b, c (Å) | 173.12 173.12 172.28 |
| α, β, γ, (° ) | 90 90 90 |
| Resolution (Å) | 34.52- 3.49 (3.62- 3.49) |
| $R_{merge}$ (%) | 0.173 (1.075) |
| I / σI | 10.67 (1.18) |
| Completeness (%) | 90.08 (51.84) |
| Redundancy | 9.6 (9.1) |
| **Refinement** | |
| No. reflections | 30527 (1723) |
| $R_{work}$ / $R_{free}$ (%) | 25.75/28.74 |
| No. atoms | |
| Protein | 8772 |
| Ligands | 4 |
| Water | 10 |
| Average B-factor | |
| Protein | 55.60 |
| Ligands | 18.61 |
| Water | 18.61 |
| R.m.s. deviations | |
| Bond lengths (Å) | |
| Bond angles (°) | 0.003 |
| Ramachandran plot statistics (%) | 0.59 |
| Most favoured | 95.23 |
| Allowed | 4.77 |
| Disallowed | 0.0 |

*Values in parentheses are for highest-resolution shell.

Values in parentheses are for the highest resolution shell. $Rmerge=\Sigma h\Sigma i|Ih,i-Ih|/\Sigma h\Sigma iIh,i$, where $Ih$ is the mean intensity of the $i$ observations of symmetry related reflections of $h$. $R=\Sigma|Fobs-Fcalc|/\Sigma Fobs$, where $Fcalc$ is the calculated protein structure factor from the atomic model (Rfree was calculated with 5% of the reflections selected randomly).

**Table S2.** SAXS experimental data statistics table of different samples

Table S2 SAXS experimental data statistics table of different samples

| | pH | Concentration mg/mL | Rg(Å) (Guinier) | I0 (Guinier) | Rg(Å) (Gnom) | I0 (Gnom) | Dmax | molecular weight（kDa） |
|---|---|---|---|---|---|---|---|---|
| SmFtsZ-WT | 7.4 | 4 | 65.5 | 421 | 65.8 | 412 | 220 | 218.4 |
| SmFtsZ-R68A | 7.4 | 4 | 53.2 | 237 | 54.7 | 235 | 188 | 166.6 |
| SmFtsZ-WT | 6.0 | 4 | 135.9 | 1240 | 137.4 | 1157 | 500 | 794 |
| SmFtsZ-R68A | 6.0 | 4 | 93.3 | 898 | 96.5 | 885 | 300 | 489 |

**Table S3.** RNA-seq results of UA159 strain and SmFtsZ-R68A strain when these strain cultured in an acidic environment.

| gene_id | UA159 | SmFtsZ-R68A | Fold Change | log2FoldChange | *p*-value | *q*-value | Regulation | description |
|---|---|---|---|---|---|---|---|---|
| SMU_RS01330 | 58.1076358 | 27.1278474 | 0.46641494 | -1.1003141 | 0.0004182 | 0.00069927 | Down | putrescine carbamoyltransferase |
| SMU_RS01335 | 27.57447371 | 18.7230753 | 0.67232089 | -0.5727781 | 0.21405343 | 0.24998566 | Down | APC family permease |
| SMU_RS01340 | 56.60291529 | 27.1317069 | 0.47899667 | -1.0619125 | 0.00080353 | 0.00130439 | Down | agmatine deiminase |
| SMU_RS01345 | 30.7145229 | 15.4371421 | 0.50398433 | -0.9885492 | 0.02054605 | 0.02811378 | Down | carbamate kinase |
| SMU_RS01325 | 111.7660616 | 43.6881021 | 0.39006324 | -1.3582201 | 1.53E-09 | 3.87E-09 | Down | helix-turn-helix transcriptional regulator |
| SMU_RS06930 | 4962.009941 | 6274.28012 | 1.26423154 | 0.33826072 | 4.15E-06 | 8.20E-06 | Up | MULTISPECIES: F0F1 ATP synthase subunit epsilon |
| SMU_RS06935 | 19943.63721 | 25030.0854 | 1.25498528 | 0.32767044 | 1.54E-05 | 2.87E-05 | Up | F0F1 ATP synthase subunit beta |
| SMU_RS06940 | 5918.904059 | 12832.4122 | 2.16760824 | 1.11610403 | 3.80E-27 | 2.61E-26 | Up | F0F1 ATP synthase subunit gamma |

| | | | | | | | | |
|---|---|---|---|---|---|---|---|---|
| SMU_RS06945 | 13448.85398 | 18230.2086 | 1.35543244 | 0.4387532 | 2.54E-05 | 4.68E-05 | Up | F0F1 ATP synthase subunit alpha |
| SMU_RS06950 | 2599.872409 | 4275.15872 | 1.64394749 | 0.71716422 | 7.21E-13 | 2.26E-12 | Up | F0F1 ATP synthase subunit delta |
| SMU_RS06955 | 2442.446433 | 4712.63715 | 1.92878035 | 0.94768886 | 2.11E-16 | 8.27E-16 | Up | F0F1 ATP synthase subunit B |
| SMU_RS06960 | 5208.440856 | 6573.15995 | 1.26182493 | 0.33551176 | 0.00536017 | 0.00795249 | Up | F0F1 ATP synthase subunit A |
| SMU_RS06965 | 3536.555905 | 1756.08159 | 0.49644955 | -1.010281 | 4.60E-09 | 1.14E-08 | Down | MULTISPECIES: F0F1 ATP synthase subunit C |
| SMU_RS07895 | 2401.540682 | 3457.36685 | 1.43903668 | 0.52510337 | 1.73E-06 | 3.53E-06 | Up | 3-hydroxyacyl-ACP dehydratase FabZ |
| SMU_RS07910 | 2346.063533 | 4135.33259 | 1.76216612 | 0.81734994 | 8.95E-17 | 3.57E-16 | Up | 3-oxoacyl-[acyl-carrier-protein] |
| SMU_RS07920 | 3638.651149 | 4108.25748 | 1.12892343 | 0.17494764 | 0.06882233 | 0.08728249 | Up | enoyl-[acyl-carrier-protein] |
| SMU_RS06135 | 353.1459037 | 398.865632 | 1.12783818 | 0.17356009 | 0.19636912 | 0.23127391 | Up | enoyl-(acyl-carrier-protein) reductase |
| SMU_RS07930 | 577.6531173 | 2970.97934 | 5.14403248 | 2.36289975 | 2.75E-137 | 8.93E-135 | Up | ketoacyl-ACP synthase III |

| | | | | | | | | |
|---|---|---|---|---|---|---|---|---|
| SMU_RS07905 | 7273.030543 | 7944.70497 | 1.09224428 | 0.12729555 | 0.22051204 | 0.25722003 | Up | beta-ketoacyl-ACP synthase II |
| SMU_RS07915 | 2976.936351 | 4840.62534 | 1.62646826 | 0.70174267 | 1.22E-16 | 4.82E-16 | Up | ACP S-malonyltransferase |
| SMU_RS07935 | 736.7678964 | 1609.38632 | 2.18175262 | 1.12548753 | 4.00E-16 | 1.54E-15 | Up | MULTISPECIES: MarR family transcriptional regulator |
| SMU_RS07940 | 3112.303236 | 7284.23437 | 2.34089873 | 1.22706253 | 7.79E-56 | 1.95E-54 | Up | enoyl-CoA hydratase |
| SMU_RS01330 | 58.1076358 | 27.1278474 | 0.46641494 | -1.1003141 | 0.0004182 | 0.00069927 | Down | putrescine carbamoyltransferase |

**Table S4.** Bacterial strains and plasmids used in this study

| Strain or plasmid | Relevant characteristic | Source or reference |
|---|---|---|
| **Strain** | | |
| *S. mutans* UA159 | Wild-type stain | Laboratory stock |
| MUT16-R68A | Arginine 68 replaced by alanine of FtsZ; Spec | This study |
| MUT17-R68L | Arginine 68 replaced by leucine of FtsZ; Spec | This study |
| MUT18-R68G | Arginine 68 replaced by glycine of FtsZ; Spec | This study |
| **Plasmids** | | |
| pET28a-smFtsZ | *E.coli* BL21(DE3), $Kana^R$ | This study |
| pET28a-smFtsZ-MUT16-R68A | *E.coli* BL21(DE3), $Kana^R$ | This study |
| pET28a-smFtsZ-MUT17-R68L | *E.coli* BL21(DE3), $Kana^R$ | This study |
| pET28a-smFtsZ-MUT18-R68G | *E.coli* BL21(DE3), $Kana^R$ | This study |
| pET28a-smFtsZ-truncated 1/319 | *E.coli* BL21(DE3), $Kana^R$ | This study |
| pET28a-smFtsZ-MUT20-Y247A | *E.coli* BL21(DE3), $Kana^R$ | This study |
| pET28a-smFtsZ-MUT19-Y247F | *E.coli* BL21(DE3), $Kana^R$ | This study |
| pET28a-smFtsZ-MUT21-E277L | *E.coli* BL21(DE3), $Kana^R$ | This study |
| pET28a-smFtsZ-MUT22-E277D | *E.coli* BL21(DE3), $Kana^R$ | This study |
| pET28a-smFtsZ-MUT23-E277A | *E.coli* BL21(DE3), $Kana^R$ | This study |
| pET28a-smFtsZ-MUT44-E277T | *E.coli* BL21(DE3), $Kana^R$ | This study |
| pET28a-smFtsZ-MUT26-T272V | *E.coli* BL21(DE3), $Kana^R$ | This study |
| pET28a-smFtsZ-MUT41-T272C | *E.coli* BL21(DE3), $Kana^R$ | This study |
| pET28a-smFtsZ-MUT25-S280A | *E.coli* BL21(DE3), $Kana^R$ | This study |
| pET28a-smFtsZ-MUT45-S280R | *E.coli* BL21(DE3), $Kana^R$ | This study |

| | | |
|---|---|---|
| pUC19-UP-smFtsZ-mNG-DN | *E.coli* Top10, $Amb^R$ | This study |
| pUC19-UP-smFtsZ_MUT16-R68A-mNG-DN | *E.coli* Top10, $Amb^R$ | This study |
| pUC19-UP-smFtsZ_MUT17-R68L-mNG-DN | *E.coli* Top10, $Amb^R$ | This study |
| pUC19-UP-smFtsZ_MUT18-R68G-mNG-DN | *E.coli* Top10, $Amb^R$ | This study |

**Table S5.** Oligonucleotide primers used in this study

| Primers | Sequence | Function |
|---|---|---|
| FtsZ_Sm-NcoⅠ-F 1 | catgCcATGGCATTTTCATTTGATGCAG | *S. mutans* FtsZ |
| FtsZ_Sm-Xho Ⅰ-R 3 | ccgCTCGAGACGATTCTTAAAGAAAGGAGG | |
| 220828 R68A-F16 | ATTAACCgcaGGTCTTGGTGCAGGAGGCCAA | pET28a-smFtsZ-MUT16-R68A/pUC19-UP-smFtsZ_MUT16-R68A-mNG-DN |
| 220828 R68A-R16 | AAGACCtgcGGTTAATTTAGGTCCAAGTTGAATAACTGT | |
| 220828 R68L-F17 | ATTAACCctgGGTCTTGGTGCAGGAGGC | pET28a-smFtsZ-MUT17-R68L/pUC19-UP-smFtsZ_MUT17-R68L-mNG-DN |
| 220828 R68L-R17 | AAGACCcagGGTTAATTTAGGTCCAAGTTGAATAACTGT | |
| 220828 R68G-F18 | ATTAACCgguGGTCTTGGTGCAGGAGGC | pET28a-smFtsZ-MUT18-R68G/pUC19-UP-smFtsZ_MUT18-R68G-mNG-DN |
| 220828 R68G-R18 | AAGACCaccGGTTAATTTAGGTCCAAGTTGAATAACTGT | |
| FtsZ_Sm-NcoⅠ-F 1 | catgCcATGGCATTTTCATTTGATGCAG | pET28a-smFtsZ-truncated 1/319 |
| smFtsZ-trca319-Xho Ⅰ-Nhis-R2 | ccgCTCGAGttaGTCTGGCCGAACACCAGT | |
| 220828 Y247A-F20 | GCGATCgcaTCACCACTTCTTGAGACGACA | pET28a-smFtsZ-MUT20-Y247A |
| 220828 Y247A-R20 | GTGGTGAtgcGATCGCCTTGCGAGC | |
| 220828 Y247F-F19 | GCGATCTtTTCACCACTTCTTGAGACGACAATTGA | pET28a-smFtsZ-MUT19-Y247F |
| 220828 Y247F-R19 | GTGGTGAAaAGATCGCCTTGCGAGCAGCCT | |

| | | |
|---|---|---|
| 220828 E277L-F21 | GAAGCTctgGAGGCTTCTGAAATTGTTAATCAAGC | pET28a-smFtsZ-MUT21-E277L |
| 220828 E277L-R21 | AGCCTCcagAGCTTCTGTCAGCGTCATATC | |
| 220828 E277D-F22 | GAAGCTgatGAGGCTTCTGAAATTGTTAATCAAGCTGC | pET28a-smFtsZ-MUT22-E277D |
| 220828 E277D-R22 | AGCCTCatcAGCTTCTGTCAGCGTCATATC | |
| 220828 E277A-F23 | GAAGCTgcaGAGGCTTCTGAAATTGTTAATCAAGCT | pET28a-smFtsZ-MUT23-E277A |
| 220828 E277A-R23 | AGCCTCtgcAGCTTCTGTCAGCGTCAT | |
| 231218 E277T-F44 | GAAGCTaccGAGGCTTCTGAAATTGTTAATCAAGCTGC | pET28a-smFtsZ-MUT44-E277T |
| 231218 E277T-R44 | AGCCTCggtAGCTTCTGTCAGCGTCATATCAAG | |
| 220828 T272V-F26 | GATATGgttCTGACAGAAGCTGAAGAGGCTTCTGA | pET28a-smFtsZ-MUT26-T272V |
| 220828 T272V-R26 | TGTCAGaacCATATCAAGGCCGCCGGTAAC | |
| 231218 T272C-F41 | GATATGtgtCTGACAGAAGCTGAAGAGGCTTCTGA | pET28a-smFtsZ-MUT41-T272C |
| 231218 T272C-R41 | TGTCAGacaCATATCAAGGCCGCCGGTAAC | |
| 221209 S280A-F25 | GAAGCTGAAGAGGCTgcaGAAATTGTTAATCAAG | pET28a-smFtsZ-MUT25-S280A |
| 220828 S280A-R25 | AATTTCtgcAGCCTCTTCAGCTTCTGTCAGC | |
| 240102 S280R-F45v2 | GAGGCTcgtGAAATTGTTAATCAAGCTGCAGGTCATGG | pET28a-smFtsZ-MUT45-S280R |
| 240102 S280R-R45v2 | AATTTCacgAGCCTCTTCAGCTTCTGTCAGC | |
| 230409 pUC19-REV-F3 | GATGCTGAAGAGATTGTTGATTTGCTGATAGAGTCGACCTGCAGGCATG | 20230409 Amplified pUC19-REV |
| 230409 pUC19-REV-R3 | AAGTGAAATCActtCACGTTCTGGTGTGATGCTCGAGGATCCCCGGGTACCGA | |
| 230409 UP-smFtsZ-F1 | TCGGTACCCGGGGATCCTCGAGCATCACACCAGAACGTGaagTGATTTCACTT | 20230409 Amplified UP-smFtsZ |

| | | |
|---|---|---|
| 230327smFtsZUp-1R | ATGCTGCATCAAATGAAAATGCCATTTTAATTTTTCCTCACTTTAATTTT | |
| 230327smFtsZ-3F | AAAATTAAAGTGAGGAAAAATTAAAATGGCATTTTCATTTGATGCAGCAT | 20230409 Amplified smFtsZ-mNG |
| 230327linker_mNG-4R | TTTATTTGCTTGTAAATCCATTAttacttgtacagctcgtccatgc | |
| 230327mNG-smFtsZDn-2F | gacgagctgtacaagtaaTAATGGATTTACAAGCAAATAAAGAACACGTT | 20230409 Amplified DN-smFtsZ |
| 230409 DN-smFtsZ-R2 | CATGCCTGCAGGTCGACTCTATCAGCAAATCAACAATCTCTTCAGCATC | |
| 230327linker_mNG-4F | TGGAAACACCTCCTTTCTTTAAGAATCGTGCAgaaGCTGCaGCtaaGgaAGC | 230403 Amplified smFtsZ-mNG |
| 230327linker_mNG-4R | TTTATTTGCTTGTAAATCCATTAttacttgtacagctcgtccatgc | |
